# Scalable Nanopore sequencing of human genomes provides a comprehensive view of haplotype-resolved variation and methylation

**DOI:** 10.1101/2023.01.12.523790

**Authors:** Mikhail Kolmogorov, Kimberley J. Billingsley, Mira Mastoras, Melissa Meredith, Jean Monlong, Ryan Lorig-Roach, Mobin Asri, Pilar Alvarez Jerez, Laksh Malik, Ramita Dewan, Xylena Reed, Rylee M. Genner, Kensuke Daida, Sairam Behera, Kishwar Shafin, Trevor Pesout, Jeshuwin Prabakaran, Paolo Carnevali, North American Brain Expression Consortium (NABEC), Jianzhi Yang, Arang Rhie, Sonja W. Scholz, Bryan J. Traynor, Karen H. Miga, Miten Jain, Winston Timp, Adam M. Phillippy, Mark Chaisson, Fritz J. Sedlazeck, Cornelis Blauwendraat, Benedict Paten

## Abstract

Long-read sequencing technologies substantially overcome the limitations of short-reads but to date have not been considered as feasible replacement at scale due to a combination of being too expensive, not scalable enough, or too error-prone. Here, we develop an efficient and scalable wet lab and computational protocol for Oxford Nanopore Technologies (ONT) long-read sequencing that seeks to provide a genuine alternative to short-reads for large-scale genomics projects. We applied our protocol to cell lines and brain tissue samples as part of a pilot project for the NIH Center for Alzheimer’s and Related Dementias (CARD). Using a single PromethION flow cell, we can detect SNPs with F1-score better than Illumina short-read sequencing. Small indel calling remains difficult within homopolymers and tandem repeats, but is comparable to Illumina calls elsewhere. Further, we can discover structural variants with F1-score comparable to state-of-the-art methods involving Pacific Biosciences HiFi sequencing and trio information (but at a lower cost and greater throughput). Using ONT-based phasing, we can then combine and phase small and structural variants at megabase scales. Our protocol also produces highly accurate, haplotype-specific methylation calls. Overall, this makes large-scale long-read sequencing projects feasible; the protocol is currently being used to sequence thousands of brain-based genomes as a part of the NIH CARD initiative. We provide the protocol and software as open-source integrated pipelines for generating phased variant calls and assemblies.

## Introduction

Most current large-scale genomics projects rely on reference mapping of short-reads to detect and genotype variants, such as single nucleotide polymorphisms (SNPs), small insertions/ deletions (indels), or structural variations (SVs) (DePristo et al., 2011). For example, short-read whole genome sequencing is routinely used in population-scale studies to discover variation in human populations (100,000 Genomes Project Pilot Investigators et al., 2021; 1000 Genomes Project Consortium et al., 2012), or to perform disease associations studies (Karczewski et al., 2020), including in cancer (Huang et al., 2018; ICGC/TCGA Pan-Cancer Analysis of Whole Genomes Consortium, 2020).

However, a substantial part of the variation in the human genome is not accessible to short reads (Sedlazeck, Lee, et al., 2018). This is because it is difficult to detect structural variants that are comparable to or longer than an individual read length (Chen et al., 2016; Mahmoud et al., 2019), resulting in missingness and error rates much higher than for small variant detection (Zarate et al., 2020; Zook et al., 2020). In addition, variation inside the repetitive regions of the genome is difficult to profile with short-reads (Wagner, Olson, Harris, Khan, et al., 2022; Wagner, Olson, Harris, McDaniel, et al., 2022) due to reference mapping ambiguity and bias (Lee & Schatz, 2012). Read-based phasing of heterozygous variants into long haplotypes is also limited by read length (Martin et al., 2016). Previous studies mostly relied on reference haplotype panels to phase known variants (Loh et al., 2016), but this method is not applicable to rare and *de novo* mutations.

Long-read sequencing, such as those from Pacific Biosciences (PacBio) or Oxford Nanopore Technologies (ONT), can overcome the limitations of short-reads and has been shown to substantially improve structural variant calling performance (Jiang et al., 2020; Sedlazeck, Rescheneder, et al., 2018) and small variant detection inside difficult-to-map parts of the genome (Shafin et al., 2021). Long-read methods can also phase small and structural variants into megabase-scale phase blocks (J.-H. Lin et al., 2022; Mahmoud et al., 2021; Shafin et al., 2021).

In addition to variant calling, several studies have used long-read sequencing to generate complete or nearly complete *de novo* genome assemblies (Logsdon et al., 2020; Rhie et al., 2021). Notably, the Telomere-to-Telomere (T2T) Consortium produced the first complete *de novo* assembly of a human genome, including centromeres and long segmental duplications (Nurk et al., 2022). Recently, the Human Pangenome Reference Consortium (HPRC) released 47 nearly-complete haplotype-resolved human genomes with diverse genetic backgrounds (Liao et al., 2022).

Despite these advances, cost and scalability have remained prohibitive barriers to the use of long-read sequencing in population-scale studies. For example, recent projects that generated high-quality genome assemblies, such as those from the Vertebrate Genomes Project (VGP), T2T, and HPRC, all used an expensive combination of multiple sequencing technologies at high coverage, including PacBio HiFi, ultra-long ONT, Hi-C, and parental sequencing (Jarvis et al., 2022).

Here, we show that it is possible to achieve state-of-the-art small and structural calling performance using only ONT reads produced by a single flow cell at a cost comparable with short-read experiments. First, we developed specialized ONT sequencing protocols that balance read length and yield. Second, we combined these sequencing protocols with a novel computational pipeline that produces haplotype-resolved de novo assemblies, along with phased small variants, structural variants, and methylation calls.

Using the data from 14 post mortem human brain samples and three human cell lines, we show that our pipeline generates SNP calls with an F1-score (harmonic mean of precision and recall) comparable to or better than state-of-the-art Illumina-based methods. Further, the structural variant calls derived from the diploid *de novo* assemblies outperform the current state-of-the-art long-read methods and are highly consistent with *de novo* assemblies produced by the HPRC consortium in genome regions outside of centromeres and segmental duplications.

Scalable long-read sequencing and associated informatics should enable the comprehensive characterization of so far inaccessible regions, including those that have been shown to harbor medically relevant mutations (Olson et al., 2022). To pursue this, we are applying the proposed methodologies across thousands of human brain samples at the Center of Alzheimer and Related Dementias (CARD) at the U.S. National Institutes of Health, with the plan to make this data openly available for research within the AnVIL cloud platform (Schatz et al., 2022). In addition, we make our sequencing and informatics pipelines openly available, the latter as a complete, easily runnable open-source software package.

## Results

### Scalable Oxford Nanopore Technologies (ONT) sequencing of cell lines and brain tissue

ONT sequencing was recently used by several studies to produce ultra-long reads (100kb+), which are ideal as an ingredient for *de novo* assembly (Jain et al., 2018; Kolmogorov et al., 2019; Koren et al., 2017; Rautiainen et al., 2022; Shafin et al., 2020). DNA preparation protocols optimized for producing very long sequence lengths typically see lower sequencing yields; therefore, multiple flow cells are used to achieve sufficient genomic coverage. In this work, we optimized a DNA processing and library preparation protocol to yield high data output (>100 Gb, corresponding to >33X coverage) in a scalable manner from a single PromethION flow-cell, while still maintaining read lengths sufficient for *de novo* assembly and long-range phasing.

The DNA processing and library preparation protocol (Methods) are publicly available through the *protocols.io* for the frontal cortex (J Billingsley et al., 2022) and cell-lines (J Billingsley, 2022). Overall, per sample, the DNA processing step yields ∼10ug of sheared DNA. Together, the DNA processing and library preparation take approximately 20 hours over two days to process up to 16 samples in a single batch, and the PromethION whole genome sequencing takes 72 hours (Figure 1).

**Figure 1.**
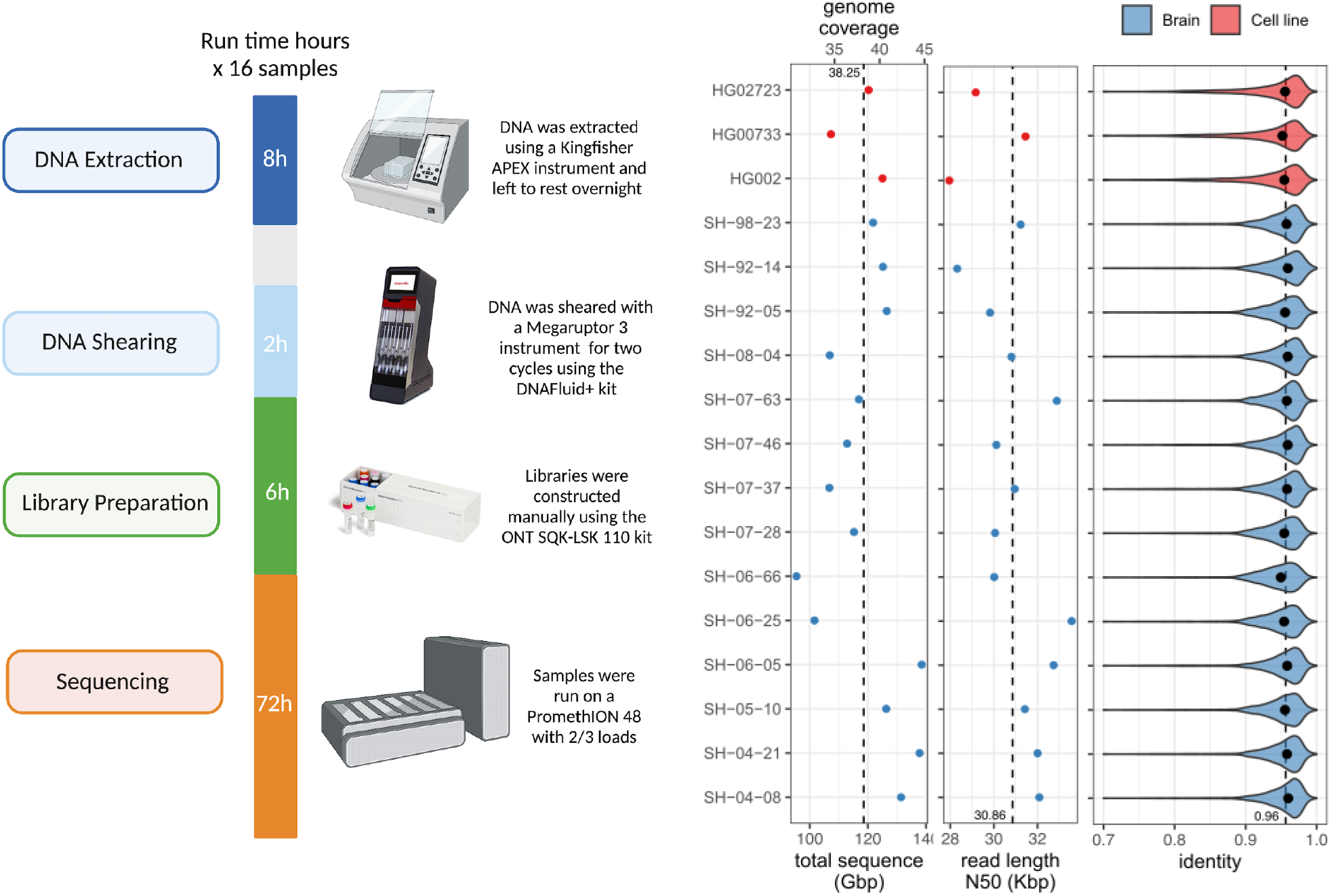
Single flow cell Oxford Nanopore Technologies (ONT) sequencing protocol. Left: Overview of the sequencing protocol, indicating all processes from DNA extraction to sequencing. In brief, the DNA is extracted using a Kingfisher Apex instrument using the Nanobind Tissue Big DNA kit. The DNA is then sheared on a Megaruptor3 instrument, and libraries are constructed using an SQK-LSK 110 kit and sequenced on a PromethION for 72 hours. Right: (from left-to-right) Total sequenced bases / haploid human genome coverage (assuming a 3.1GB genome) from PASS reads (with estimated QV>=10) for each sample. The vertical dotted line marks the average yield across samples. Read length N50 of PASS reads, i.e., the read length (y-axis) such that reads of this length or longer represent 50% of the total sequence. The vertical dotted line marks the average N50 across samples. Distribution of PASS read identities when aligned to T2T-CHM13 v2.0. The dots mark the median identity in each sample, and the vertical dotted line is the average across samples.

Using our optimized protocol, we sequenced 17 human genomes, including three cell lines (HG002, HG00733, HG02723) that have been extensively used in other benchmarking studies, and 14 brain tissue samples that were obtained as a part of NABEC cohort (Gibbs et al., 2010). Each dataset was sequenced using a single PromethION flow cell, yielding on average 116 Gb of base-called reads (∼37X coverage assuming 3.1Gb genome) with an estimated average across bases read quality above Q10. The average read N50 was 31 kb (average maximum = 769 kb), which is lower overall than other long DNA preparation protocols (Shafin et al., 2020). However, there is a substantial reduction in cost, improved scalability, and high consistency (Figure 1; Supplementary table 1). We used the R9.4.1 pore version and the Guppy version 6.1.2 in super accuracy mode for base and methylation calls, median read identity mapped to GRCh38 was 96%.

### Variant calling and methylation analysis pipeline overview

We adapted existing tools and developed new methods for high-quality variant calling, haplotype-specific methylation profiling, and diploid *de novo* assembly (Methods; Supplementary Figure 1). The pipeline begins by generating a diploid *de novo* assembly using a combination of Shasta, which produces a haploid assembly and Hapdup (https://github.com/KolmogorovLab/hapdup), which generates locally phased diploid contigs. We then use the generated assemblies to call structural variants (at least 50 bp in size) against a given reference genome using a new assembly-to-reference pipeline called hapdiff (https://github.com/KolmogorovLab/hapdiff; Methods).

Ideally, small variants could also be recovered from diploid contigs, as has been successfully done for HiFi-based assemblies (Liao et al., 2022; Wagner, Olson, Harris, McDaniel, et al., 2022). Our Shasta-Hapdup assemblies had mean substitution error rates of ∼8 per 100 kb, which is similar or better compared to the previous ONT-based assemblies (Kolmogorov et al., 2019; Shafin et al., 2020, 2021), but higher than current contig assemblies produced with PacBio HiFi (<1 per 100kb) (Liao et al., 2022). Reference-based small variant calling methods are less error-prone because they can better approximate the biases of ONT base calling errors via supervised learning (Shafin et al., 2021). Thus, in our current pipeline we used an updated version of the PEPPER-Margin-DeepVariant software (Shafin et al., 2021) to call small variants against a reference.

Given a set of structural variants produced with *de novo* assemblies, and reference-based small variant calls, our pipeline phases them into a harmonized variant call set using a newly introduced functionality of the Margin tool. In addition, given the phased reference alignment with methylation tags (produced by Guppy), we produce haplotype-specific calls of hypo- and hyper-methylated regions of the genome.

We publish the complete open-source pipeline written in WDL in the Dockstore repository (https://dockstore.org/organizations/NIHCARD; Methods) to encourage easy reuse. Our analysis was run on the cloud using the Terra compute environment (Schatz et al., 2022). Analysis of a single ONT human sample at 30-40x coverage took 22-25 wall-clock hours if run on the Terra environment (2200-2500 CPU hours; estimated Google Cloud computing cost about 100$; Supplementary Table 1).

### Small variant calling and benchmarking

We benchmarked the performance of small variant (substitutions and indels shorter than 50 bp) calls produced by the updated PEPPER-Margin-DeepVariant and compared them to Illumina-based calls produced by DeepVariant (Figure 2; Supplementary Table 2). Comparisons are reported using false positives (FP) and false negative (FN) and F1 score (harmonic mean of precision and recall) metrics. First, we used the Genome in a Bottle (GIAB) small variant benchmark (v4.2.1) that provides a curated set of small variant calls for the HG002 genome. Inside the Tier1-confident regions, both ONT and Illumina-based SNP calls were highly concordant with the GIAB benchmark (F1-scores 0.997 and 0.996, respectively). The residual error was however noticeably lower for ONT-based SNP calls (FP + FN = 15,916), compared to Illumina-based calls (FP + FN = 23,203).

**Figure 2.**
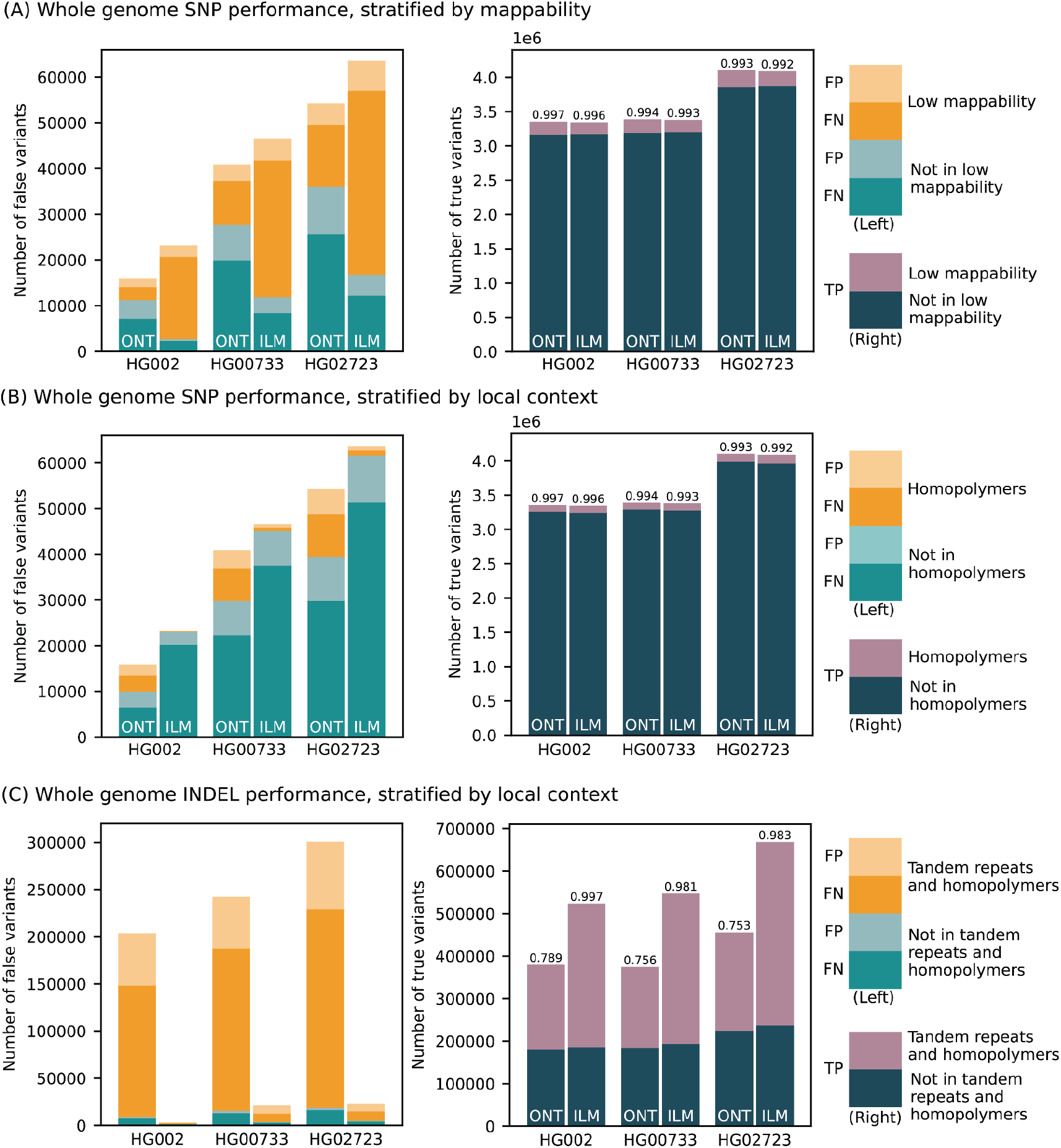
Small variant calling performance evaluation. (Left panels) Number of false positive (light bars) and false negative (dark bars) calls made by PEPPER-Margin-DeepVariant (PMDV) using ONT reads and DeepVariant using Illumina reads. Statistics computed against the Genome in a Bottle v4.2.1 benchmark for HG002; for other cell lines (HG00733, HG02723) calls generated by DeepVariant with HiFi reads are used. Whole genome SNP counts are stratified by mappability in (A) and local context in (B); INDEL counts by local context in (C). (Right panels) Number of true positive variant calls stratified by different genomic intervals. F1-score is reported on top of each bar.

Since the GIAB benchmark is only available for the HG002 genome, we used small variant calls from HiFi reads (produced by DeepVariant) as ground truth for the HG00733 and HG02723 genomes. The results were consistent with the GIAB benchmark analysis, confirming the reduced SNP error rate of ONT-based methods, compared to Illumina within the Tier1 benchmark regions (Figure 2; Supplementary Table 2).

The difference in recall and precision between ONT and Illumina-based calls was dependent on the genomic context (Figure 2). The major source of error in Illumina-based calls is due to false negatives within regions of known low mappability. Inside those regions, ONT-based calls had noticeably higher SNP F1-score (0.987) compared to Illumina (0.944). Conversely, Illumina performance was better in homopolymers of size at least 7bp (SNP F1-score was 0.999 for Illumina and 0.970 for ONT) and, to a lesser extent, within tandem repeats (SNP F1-score 0.997 for Illumina and 0.992 for ONT).

Small indels continue to be the major source of error for ONT-based variant calls (F1 score of 0.789 for ONT vs. 0.997 for Illumina inside the Tier1 GIAB regions) (Figure 2). The large majority of indel errors in ONT calls occur within homopolymers and tandem duplications (F1 score of 0.676 for ONT vs. 0.997 for Illumina). Outside of homopolymer runs and tandem repeats (approximately 35% of all indels), F1-scores for both ONT and Illumina were much more comparable (F1-score 0.975 and 0.996, respectively). In exons (defined by the GIAB benchmark and representing <0.1% of all indels) F1-scores were also more comparable (0.9230 for ONT and 0.9923 for Illumina; Supplementary Table 2). In addition, our analysis of R10 sequencing of the cell lines shown below highlights substantial improvements in the accuracy of small indel calls.

### *De novo* haplotype-resolved assembly using only ONT reads

Recent studies utilized HiFi-based *de novo* assemblies to produce highly accurate and complete structural variant calls (Liao et al., 2022; Wagner, Olson, Harris, McDaniel, et al., 2022). To profile the full spectrum of heterozygous variants, diploid assembly is required; HiFi-based studies used either trio or Hi-C information to produce phased assemblies. Although Shasta has a somewhat experimental phased assembly mode designed to work with ultra-long (>= 100 kb) reads, the default modes of current ONT assembly methods (such as Shasta or Flye) only generate haploid assemblies, representing a random mosaic of the paternal and maternal haplotypes. We therefore developed a pipeline called Hapdup that takes a haploid assembly as input and produces a locally phased diploid assembly, sometimes also referred to as a dual assembly (Methods).

Using the Shasta + Hapdup combination, we generated *de novo* assemblies for 14 brain samples and three cell lines (Figure 3; Supplementary Table 3). All assembly statistics were highly consistent among the samples, with the mean haploid assembly size of 2.88 Gb; mean NG50=22.0 Mb; mean NGA50=14.6 Mb (measured against the T2T-CHM13 2.0 reference). Mean QV (34.22) was estimated from k-mer frequencies using yak. Mean NG50 of the phase blocks was ∼2 Mb; switch error rate of cell line assemblies was 0.07-0.18% (Supplementary Figure 2).

**Figure 3.**
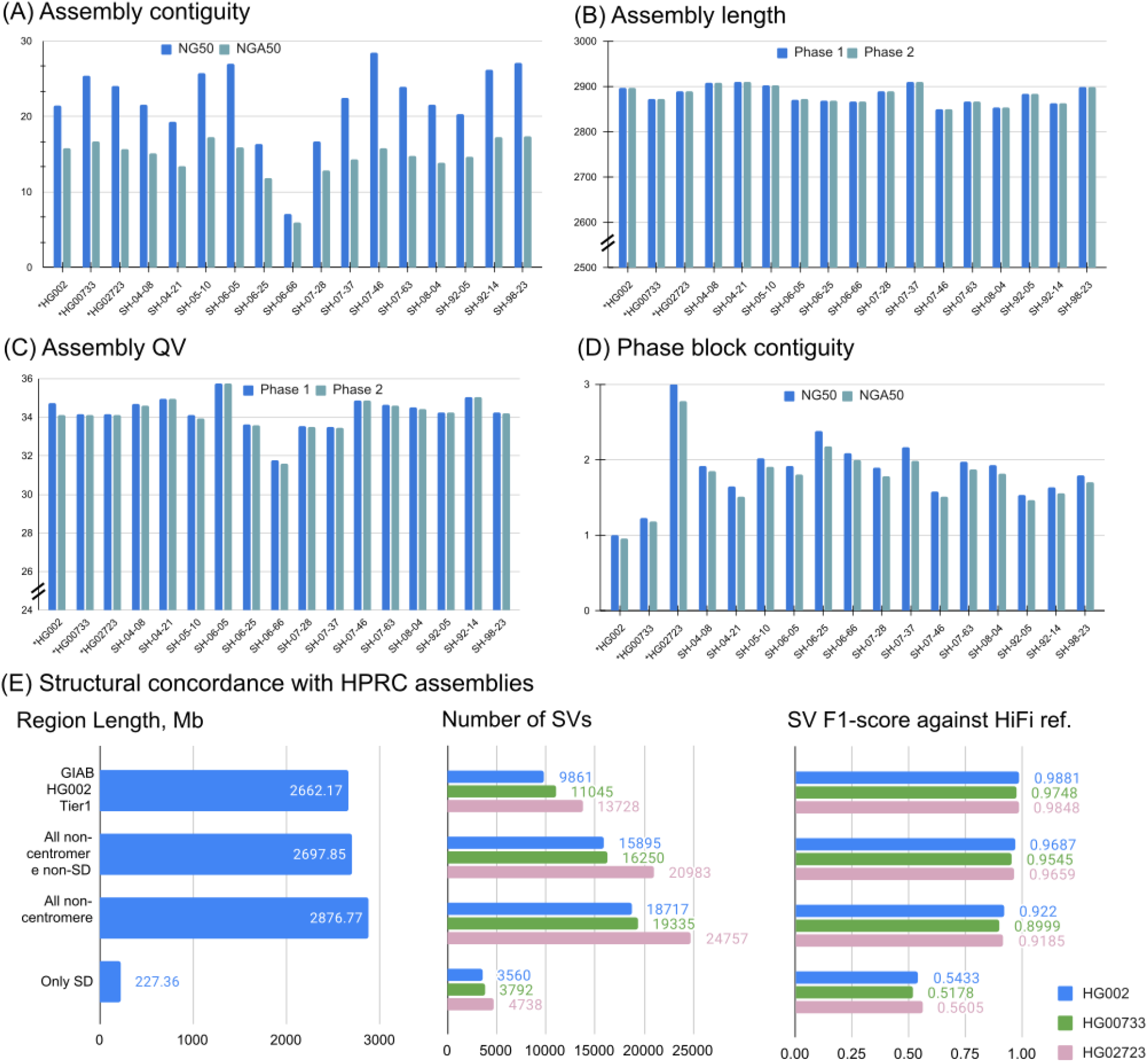
Assemblies of 14 brain tissues and 3 cell lines generated by Shasta+Hapdup. (A) NG50 and NGA50 contiguity measured using QUAST. Sample 06_66 had the lowest contiguity due to the decreased sequencing yield. (B) Assembly length. (C) Mean assemblies QV computed using yak. (D) Contiguity of phased blocks, broken at phase switches. An increased value for HG02723 suggests an increased heterozygosity rate. (A-D) Cell lines marked with asterisks. (E) Structural variation call concordance with HiFi-based assemblies for various regions of the genome.

Compared to the trio-binned ONT HG002 assemblies produced from a recent long-read benchmarking study (Jarvis et al., 2022), our assemblies had comparable contiguity and better QV and phasing accuracy (Supplementary Figure 3). Compared to trio-binned HiFi HG002 assemblies, our assemblies had lower QV due to the residual error in ONT-based consensus.

We used Flagger (Liao et al., 2022) to detect potential errors in Shasta+Hapdup assemblies (Methods; Supplementary Figures 12 and 13). Outside of centromeres and segmental duplications, 0.69 - 1.98% of the assembled sequence was marked as unreliable by Flagger (for autosomal contigs longer than 1 Mb; using HiFi reads for validation). In comparison, 0.12 - 0.27% was reported in the assemblies released by the HPRC (which integrated multiple sequencing technologies including PacBio HiFi) - a lower, but comparable fraction. This confirms high structural accuracy of our assemblies outside of major repeats, in agreement with the structural variant analysis described below (Figure 3e). Segmental duplication assemblies produced by Hapdup were less reliable, primarily due to the reduced read length of our sequencing protocol (23.33 - 25.12% of segmental duplication sequence was reported unreliable with HiFi reads; Supplementary Figures 12 and 13).

### Structural variant calling and benchmarking

To produce assembly-based SV calls, we developed a package called hapdiff, which is a combination of minimap2 and a modified version of SVIM-asm (Methods). In particular, we added functionality to group multiple indels inside a single VNTR element.

We benchmarked Hapdup and HPRC assembly-based SV calls, as well as reference mapping-based SV calls produced with Sniffles2 (Smolka et al., 2022) and CuteSV (Jiang et al., 2020). We also included a comparison with short-read-based SV calls using Manta, one of the top performers in the recent short-read SV studies (Zarate et al., 2020; Zook et al., 2020). We benchmarked these sets of SV calls using Truvari (English et al., 2022) against the recent manually curated structural variant benchmark produced by the Genome in a Bottle initiative for the HG002 genome (Zook et al., 2020).

Hapdup-based SV calls (F1-score 0.967) improved over Sniffles2 (0.953) and CuteSV (0.938) that were also run using the ONT data. HPRC-based assembly concordance was only slightly higher (0.970), compared to Hapdup (Figure 4; Supplementary Table 4). Illumina-based SV calls had substantially lower performance (F1-score 0.402), in particular missing many insertions (Supplementary Figure 4). In addition, long insertions are often misclassified as translocations by short-read methods (Sedlazeck, Rescheneder, et al., 2018).

**Figure 4.**
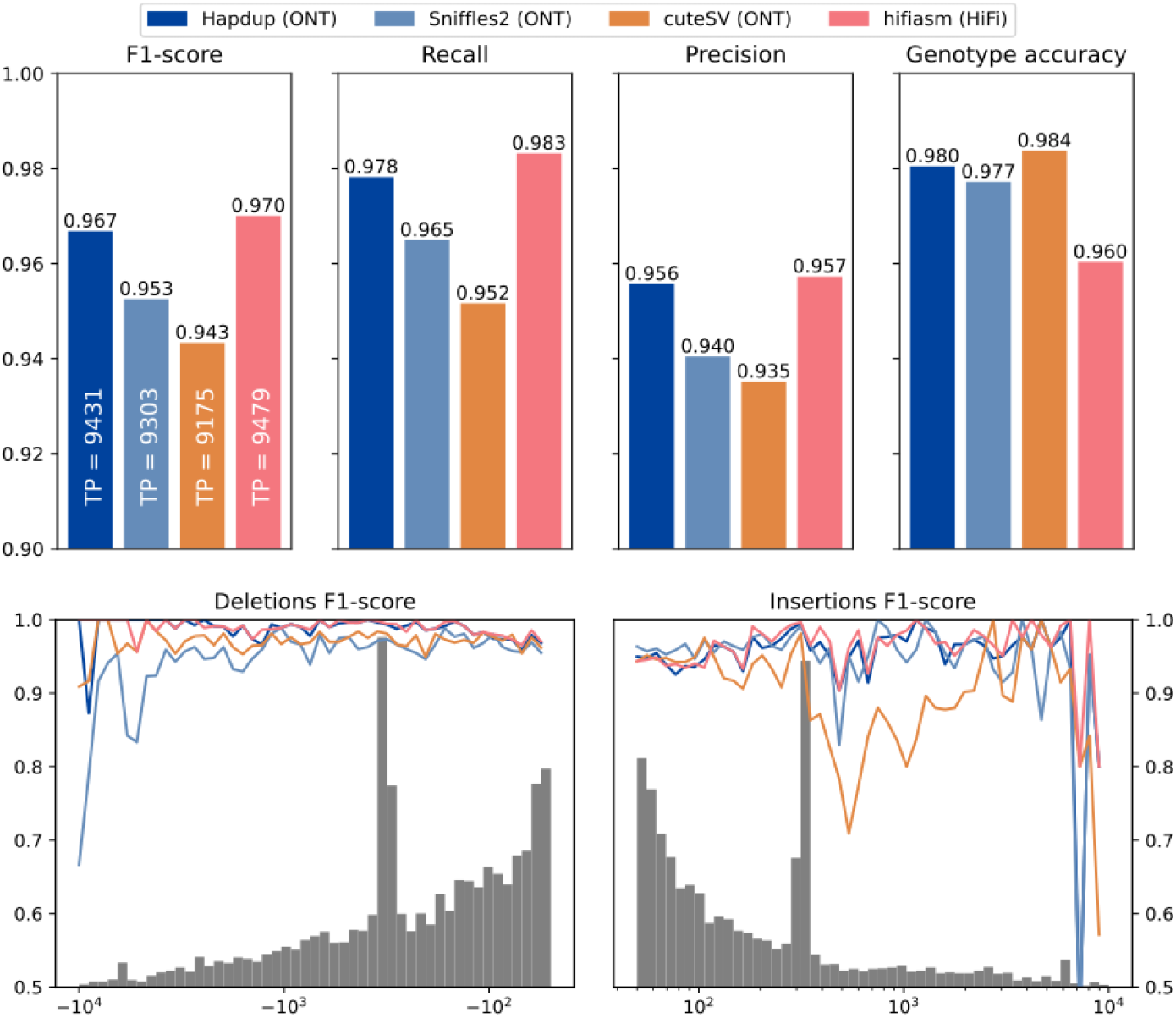
Structural variant evaluations using the GIAB HG002 benchmark. (Top) Recall, precision and F1-scores computed for various tools and sequencing technologies with Genome in a Bottle Tier1 v0.6 benchmark as reference (defined on HG002). (Bottom) F1-score computed for various SV size bins. The gray histogram shows the distribution of SV sizes in the reference set.

The GIAB call set is limited to a set of high-confidence regions and lacks variant calls in many complex loci. To investigate the quality of our SV calls further, we used the assemblies generated by the HPRC as a benchmark on three cell lines and evaluated the concordance of SV calls inside various regions of the human genome. While the HPRC assemblies are not perfect, they do span nearly the entire genome. Using the T2T-CHM13 assembly as a reference, we filtered out annotated centromere satellite repeats and segmental duplications. The remaining regions contain 15,588 SVs in HG002 HPRC-based calls, a ∼50% increase compared to the GIAB SV (v0.6) Tier1 regions. Hapdup retained a high agreement with HiFi assemblies on these regions (F1-score 0.95-0.97). In the regions with only satellite repeats filtered out, therefore keeping segmental duplications, Hapdup had a reduced F1-score of 0.93, and these regions contained 16,219 SVs total. Analysis with two other cell lines showed similar trends (Figure 2e).

Supplementary Figure 5 Illustrates that using VNTR SV grouping improves the concordance between SV sets produced by Hapdup and HPRC assemblies. Sniffles2 had comparable F1-score against the HPRC-based SV calls inside GIAB Tier1 regions, but the concordance was substantially reduced for the extended genomic intervals relative to the Shasta/HapDup calls (Supplementary Figure 6). This may partly be explained by the ambiguities in SV representation rather than truly erroneous calls, in particular representation of indels inside VNTRs. Analysis using the TT-Mars tool that performs SV calls validation on the sequence level also confirmed the high quality of Hapdup-based SV calls relative to other tools (Supplementary Figure 7).

Evaluations are also shown for a subset of SNPs that are within 100 bp of structural variants (Bottom) An example of a Hapdup and hifiasm representations of complex clusters with small and structural variants at chr1:55,544,500-55,551,000 (in CHM13 reference), visualized using IGV. Top tracks show phased SNPs and SVs produced by our pipeline and derived from HPRC assemblies (using dipcall). A few inconsistencies between SNP positions are explained by ambiguities between read and contig alignments around SV sites.

### Improved representation of variants in challenging medically-relevant loci

We further assessed the ability of our assemblies to represent and cover medically challenging genes (Wagner, Olson, Harris, Khan, et al., 2022; Wagner, Olson, Harris, McDaniel, et al., 2022). Here we assessed 389 genes that we previously postulated as highly complex and repetitive (e.g., *LPA*, *SMN1*, *SMN2*). We first measured the number of contigs observed per region to identify the coverage, and we observed that 359 and 356 genes are covered by at least one contig from the first haplotype and second haplotype, respectively (see Supplementary Table 4). Given that we cover these medically challenging genes, we next assessed the variant calling ability in these genes. For this, we used the recent benchmark from GIAB (CMRG v1.0) for SNPs and SVs. We were particularly interested in the ability of our method to recover indels and SVs. For substitutions, our reference-based pipeline obtained a high accuracy (F1-score: 0.97). For smaller insertions and deletions (<50bp), we obtained an accuracy of 0.68 in these repetitive challenging genes. Next, we assessed the SV performance. Given the limited set of genes in this benchmark, we only could assess the performance based on 252 SVs. Our assemblies achieved a high accuracy (F1-score: 0.91); notably, SV calling using hapdiff achieved higher F1-score compared to SV calls made using dipcall (F1-score: 0.88). Sniffles2 had a similar F1-score on this benchmark (0.91; Supplementary Table 4).

### Harmonized phased representation of small and structural variants

Structural and small variant calls were merged using bcftools concat, then harmonized and phased with Margin to produce a complete representation of the sample variants (Methods). Mean phase block NG50 was 1.08 Mb among the brain samples (Figure 5). HG002 and HG0073 cell lines had similar statistics, but HG02723 had noticeably higher phase block length (NG50 2.83 Mb). This individual has a higher proportion of African ancestry, and as a result, fewer apparent blocks of autozygosity that prevent phasing.

**Figure 5.**
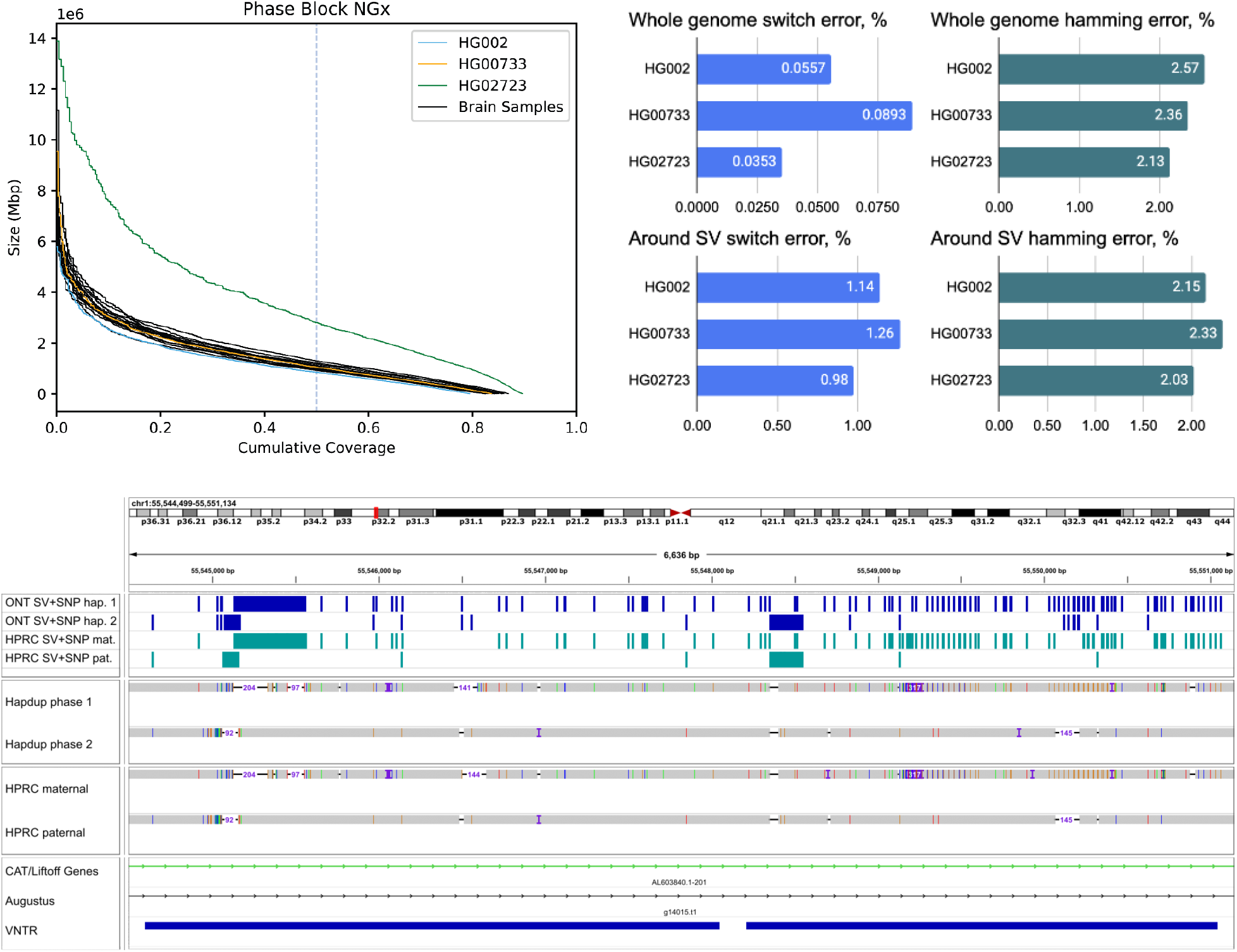
Combined, phased small and structural variants improve the profiling of complex genomic regions. (Top) Variant phasing evaluation. Left plot shows the phased block NGx, reported by Margin. HG02723 has an increased phase block length due to higher heterozygosity. Right plots show SNP hamming and switch error computed against the small variants in HiFi-based assemblies.

We evaluated phasing accuracy of the cell line calls against the small variant calls produced from HPRC-based assemblies using the “whatshap compare” module. Switch error (ratio of adjacent SNPs in a wrong phase) was 0.04-0.09%, comparable to assemblies produced with a combination of HiFi and Trio / Hi-C (Cheng et al., 2021, 2022). Hamming error (ratio of all SNPs inside a phased block in a wrong phase) was also low (2.1-2.6%), albeit slightly higher compared to Trio / Hi-C-based assemblies.

Notably, the original phasing approach that only considers small variants had substantially higher switch error rates (0.17-0.21%; Supplementary Table 2). This highlights that considering SVs improves phasing quality, consistent with other recent studies (J.-H. Lin et al., 2022). The switch error around SVs (within 100 bp from boundaries) was 1-1.26%, which was an improvement over 2-2.4% rate of the SV-unaware method (Figure 5; Supplementary Table 2), but nevertheless elevated compared to the rest of the genome. More than half of SVs coincide with difficult-to-map VNTR regions, which may explain the reduced performance.

A harmonized, phased view of all variants can facilitate the characterization of complex regions that contain multiple small and structural variants on both haplotypes. Figure 5 presents an example of such a region in the HG002 genome; our variant representation of this region provides an integrated view of variation that is consistent with the HPRC assemblies. These cases are not rare; overall, our assemblies had 3,149-5,554 SVs co-located with other SVs (Supplementary Figure 8).

### Advantages of using the T2T-CHM13 reference genome

In our evaluations, we used both GRCh38 and T2T-CHM13 references to call small and structural variants. T2T-CHM13 contains approximately 200 Mb of sequence that is missing, misrepresented, or simulated in GRCh38, allowing the mapping and calling of more variants, particularly in sequences rich in repetitive content. Reflecting this, in three cell lines, PEPPER-Margin-DeepVariant called 1.19 - 1.29 million variants in T2T-CHM13 sequence non-syntenic to GRCh38, compared to 0.26-0.28 million variants in GRCh38 sequence non-syntenic to T2T-CHM13 (Supplementary Figure 9).

To evaluate the small variant consistency, we lifted over GIAB confident regions from GRCh38 to T2T-CHM13. Our ONT-based small variant calls had high F1-score similarity (0.9951) with the calls produced using DeepVariant and HiFi data against the T2T-CHM13 reference. The F1 similarity was slightly below the concordance between ONT calls and HiFi calls using the GRCh38 reference (F1-score 0.9976), which is likely explained by a few liftover artifacts (Supplementary table 2).

### Analysis of rare structural variants in brain genomes

We next analyzed the distribution of structural variants in the 14 brain samples, which have not been previously sequenced with long-reads. The analyzed samples represent control cases from the NABEC cohort (age range, 68 - 95 years), which at the time of death did not show signs of neurodegenerative disorders.

On average, 19,255 SVs were identified in the most confident regions of the genome (Figure 6A; Supplementary Figure 10; Supplementary Table 6). The SVs were matched to three SV catalogs to annotate their frequency in the population. In each sample, about 304 (1.6%) SVs were absent from the public SV catalogs or matched with rare variants. Among those rare variants, about 14 per sample are located around genes, including ∼4 on average overlapping coding regions (Figure 6A, right).

**Figure 6.**
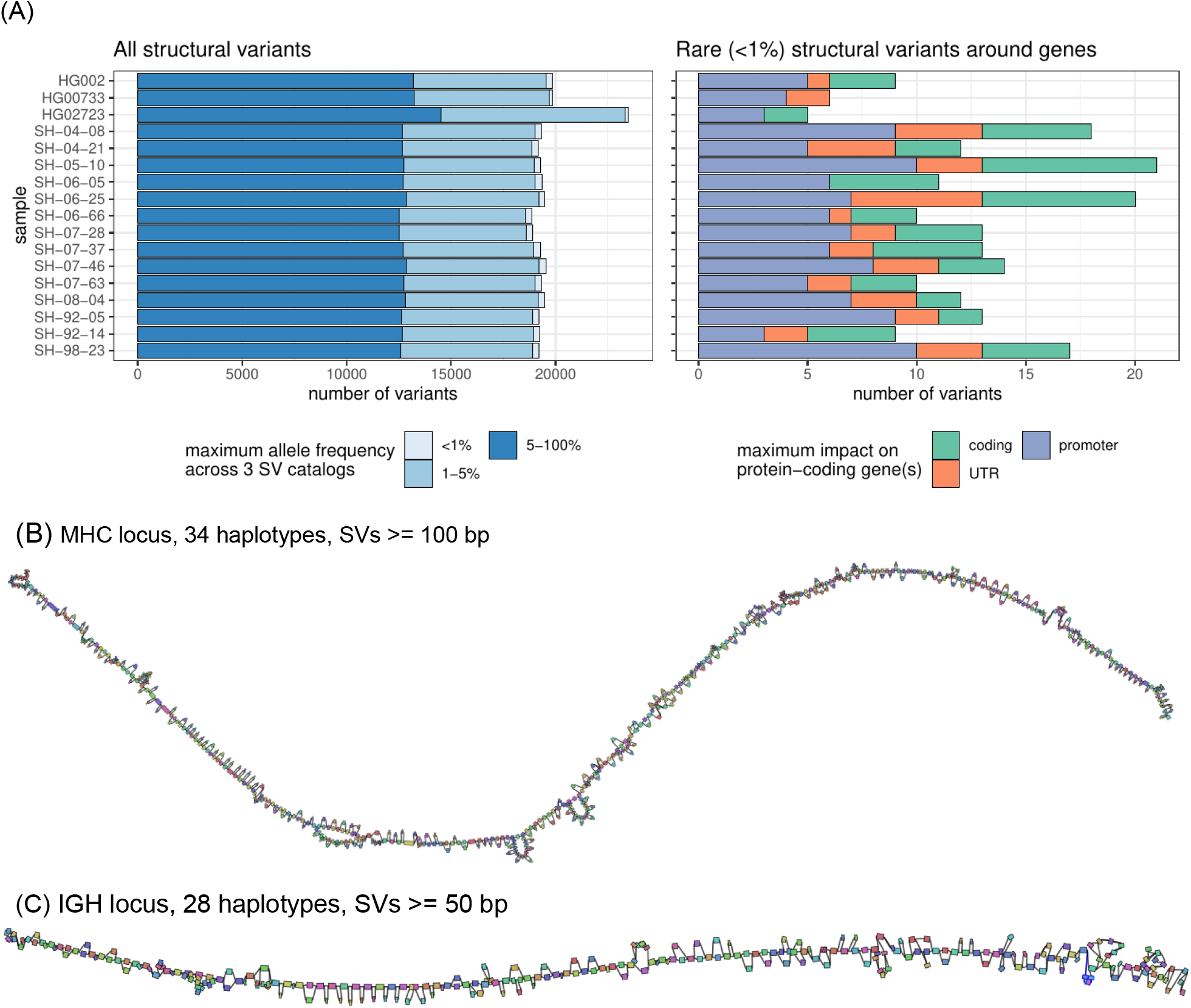
Structural variant landscape summary. (A) The number of structural variants across samples. In the left panel, structural variants were annotated with three SV catalogs (the gnomAD-SV database, a long-read-based SV catalog, and the HPRC v1.0 SV catalog). SVs are matched if they have at least 10% genomic overlap. SVs close to centromeres, telomeres, or within segmental duplications were removed. The colors highlight the maximum frequency across these catalogs, the lighter blue showing “rare” SVs (with an allele frequency below 1%) in the catalogs, or unmatched. SVs may be unmatched, either because they are novel or due to the difficulties in the database comparison. The right panel shows the number of rare structural variants in protein-coding genes, grouped by their impact on the gene structure. (B) MHC pangenome built from 28 brain and 6 cell line haplotypes, containing 640 nodes, SVs over 100bp are shown. (C) IGH pangenome built from 28 brain haplotypes containing 268 nodes. In contrast, cell lines are typically derived from B-cell lymphocytes and contain extensive somatic rearrangements in this locus.

Despite being control samples, we found 3 SVs in total amongst the SVs coinciding with coding regions that overlapped coding sequences of genes that are predicted to be intolerant to loss-of-function variants or to be haplo-insufficient. One such example is a 4.2 Kbp heterozygous deletion of a transcription start site and exon of *RBFOX1*, a gene that may be involved in spinocerebellar ataxia type 2, a rare neurological condition (Supplementary Figure 11).

Our brain genome assemblies contained contiguous assemblies of highly polymorphic loci. To investigate such loci, we looked at the major histocompatibility complex (MHC) and immunoglobulin heavy chain (IGH) loci, which were both assembled in single haplotype-specific contigs in all our samples. Reference-based characterization of these regions is difficult because of high heterozygosity and repetitiveness. We instead constructed pangenome graphs (using minigraph) that represent all structural variants (Liao et al., 2022). The MHC pangenome graph was built from 28 brain and 6 cell line haplotypes and consists of 1,294 nodes reflecting SVs larger than 50 bp; we also built a “lower resolution” graph with 640 nodes containing SVs over 100 bp (Figure 6B). An IGH pangenome was built from 28 brain haplotypes and contained 268 nodes with SVs larger than 50 bp (Figure 6C). Cell lines are typically derived from B-cell lymphocytes and contain extensive somatic rearrangements in this locus and are therefore not suitable for the germline variant comparison.

### Haplotype-resolved methylation calls

ONT sequencing also allows the identification of base modifications. Here, we produced phased 5-methylcytosine calls aligned to the GRCh38 and T2T-CHM13 references. Initial methylation calls were produced *de novo* using Remora (https://github.com/nanoporetech/remora).

For the HG002 genome, our methylation calls covered 28.83 million CpG sites (98.8% of total GRCh38 CpG sites) and had a high correlation to calls made using the standard whole genome bisulfite sequencing (WGBS) in regions covered by both technologies (R=0.949, RMSE=11.314; Figure 7a). We calculated correlations between all other samples and HG002 WGBS data to understand the level of ‘background’ methylation between all samples; these correlations ranged between 0.84-0.86 (Supplementary Table 5). HG002 WGBS calls were collected from the ONT open data repository (https://labs.epi2me.io/gm24385-5mc).

**Figure 7.**
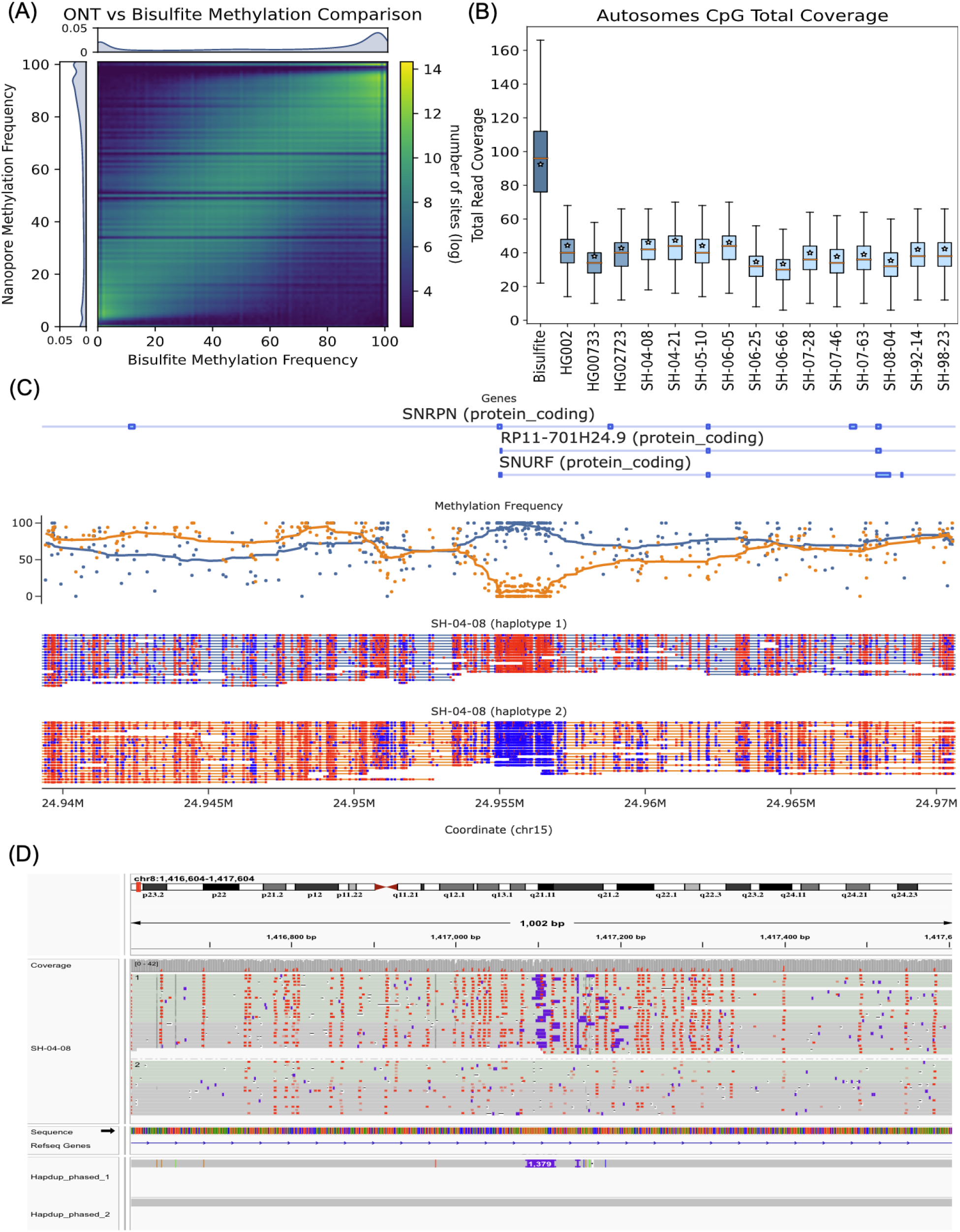
Haplotype-specific methylation profiling. (A) Heatmap of concordance between Bisulfite whole genome sequencing and ONT Remora Methylation calls in HG002 at sites shared by both technologies covered by at least 5 reads. The lower coverage of ONT data causes striping in the heatmap at specific frequencies. (B) Read depth of Bisulfite and ONT samples, this plot shows that less ONT coverage is able to obtain the same methylation information as bisulfite with more than twice the coverage. CpG sites are one position apart in the sense and antisense DNA strands due to C-G base pairing. Since this read coverage is counted per CpG location the actual coverage was doubled to account for the neighboring strand locations and estimate actual genome wide coverage. (C) A positive control plot showing the expected differential methylation pattern in the *SNRPN* (Small Nuclear Ribonucleoprotein Polypeptide N) of phased ONT reads for brain sample SH-04-08. Red CpG sites are methylated and blue sites are unmethylated. Above the reads is a plot of methylation frequency and gene locations, visualized using modbamtools (D) IGV visualization of phased methylated ONT reads and the phased assemblies of brain sample SH-04-08 at the gene DLGAP2 locus that shows a 1,379 base pair insertion that is differentially methylated across haplotypes.

A unique feature of ONT-based methylation calls, compared to WGBS is haplotype-level resolution (Figure 7c). To explore this, we identified haplotype-specific differentially methylated regions in gene promoter regions and regions flanking structural variants. Here we consider gene promoters that have a difference in average methylation between haplotypes that are more than three deviations away from the absolute median difference.

Differential haplotype methylation patterns were found in 4.73% (690) of autosomal protein-coding gene promoters in HG002. Similarly, HG00733 and HG02723 had 4.43% (662) and 3.57% (519) gene promoters differentially methylated, respectively. The brain samples had 2.8% - 3.8% of promoters differentially methylated. Differential haplotype methylation for the UCSC GRCh38 CpG islands track ranged from 6% - 7.6% (1651 - 2109) across brains and cell lines. Structural variants also showed differential haplotype methylation patterns in 5.7% - 6.2% (532 - 731) of deletions in cell lines and 4.6%-5.8% (454 −554) of deletions in brain samples. Figure 7D shows an example of gene DLG associated protein 2 (DLGAP2) in SH-04-08 sample with a 1,379 bp heterozygous insertion that coincides with haplotype-specific methylation.

### Comparing nanopore sequencing of cell lines using R9 and R10 pore versions

During the preparation of this manuscript, a newer version of the nanopore sequencing protocol (R10.4.1 kit V14) became commercially available. To evaluate the benefits of the newer protocol, we re-sequenced HG002, HG00733 and HG02723 cell lines and compared the results with the R9 protocol. Using sample preparation procedures similar to the described above, we generated 110-121 Gb of reads with N50 ∼29 kb using single flow cells (Table 1). The median read identity was 99%, an improvement compared to 96% using R9 sequencing. We slightly modified the computational pipeline by providing a specialized R10 model for DeepVariant and configuration file for Shasta, but no other changes were required.

**Table 1.** Sequencing statistics of PASS reads from R9 and R10 Kit V14 sequencing of three cell lines using single PromethION flow cells.

| Dataset | Total yield (Gb) | Coverage | Read N50 (kb) | Median GRCh38 identity |
| --- | --- | --- | --- | --- |
| HG002 R9.4.1 | 125.0 | 40.2 | 27.9 | 0.9544 |
| HG002 R10 Kit V14 | 121.0 | 38.9 | 29.4 | 0.9879 |
| HG00733 R9.4.1 | 120.2 | 38.7 | 29.1 | 0.9518 |
| HG00733 R10 Kit V14 | 110.0 | 35.4 | 30.8 | 0.9866 |
| HG02723 R9.4.1 | 107.3 | 34.5 | 31.4 | 0.9518 |
| HG02723 R10 Kit V14 | 113.2 | 36.4 | 32.2 | 0.9869 |

Assemblies and variant calls using R10 showed better or similar performance in almost all benchmark categories (Table 2). Notably, Shasta+Hapdup assembly QV improved from 34.3 to 42.8, resulting in substantial improvements in assembly-based SNP calls (recall and precision >0.985). The improved assembly quality also resulted in slight improvements in SV calls against GIAB benchmark and HPRC calls. Although HG002 assembly NG50 decreased from 21.4 Mb to 14.9 Mb, this had no effect on the structural variant calling performance. Further optimizations in Shasta assembly parameters will likely improve R10 assemblies contiguity.

**Table 2.** Comparison of R9 and R10 sequencing of the HG002 cell line. Substantial improvements in different categories are highlighted in bold. Benchmarks were performed using the methods described above in the manuscript.

| Benchmark | HG002 R9 | HG002 R10 kit V14 |
| --- | --- | --- |
| Assembly benchmarks |  |  |
| Asm. size | 2.896 Gb | **2.927** Gb |
| Asm. NG50 | **21.4 Mb** | 14.9 Mb |
| Asm. phase block N50 | 1.010 Mb | 0.99 Mb |
| Asm. SNP switch | 0.00165 | 0.0015 |
| Asm. QV | 34.3 | **42.8** |
| Asm. SNP recall / precision | 0.9795 / 0.9528 | **0.9851 / 0.9856** |
| Small variant calls benchmarks (recall / precision) |  |  |
| SNP (GIAB Tier 1) | 0.997 / 0.9982 | 0.9979 / 0.998 |
| Indel (GIAB Tier 1) | 0.7217 / 0.8715 | **0.8495 / 0.8991** |
| Indel (no homopol. or tandem repeats) | 0.9609 / 0.9807 | **0.9970 / 0.9969** |
| Indel (RefSeq CDS) | 0.9121 / 0.9342 | **0.9948 / 0.9748** |
| Structural variant benchmarks (recall / precision) |  |  |
| SV (GIAB Tier1) | 0.9782 / 0.9557 | 0.975 / 0.9595 |
| SV (HPRC non-Cen non-SD) | 0.9689 / 0.9685 | **0.9764 / 0.9835** |
| SV (HPRC only SD) | 0.4921 / 0.6064 | **0.5277 / 0.6355** |

As expected, the R10 protocol resulted in substantial improvements in reference-based indel calls. Inside GIAB Tier1 regions, F1-score improved from 0.79 to 0.87 (R9 and R10, respectively). Notably, outside of homoploymers and tandem repeats, R10 indel calls also improved substantially and were similar to Illumina calls (F1-scores 0.997 and 0.996, respectively). Similarly, indel F1-scores in exons improved from 0.923 to 0.985 (R9 and R10, respectively). Reference-based SNP calls were comparable between R9 and R10.

Overall, the R10 pore version represents a clear improvement over R9 and we expect the CARD initiative to use the R10 pore for sequencing of new cohorts. Notably, our wet-lab and computational methods required little adjustments and R9 and R10 benchmarks were overall highly concordant. The specialized R10 version of the WDL workflow is available in the “r10” branch of the main repository.

## Discussion

In this work, we designed an ONT sequencing protocol that produces over 100Gb of ONT reads using a single PromethION flow cell. We developed new workflows and adapted and updated existing tools, combined into an end-to-end pipeline implemented in WDL, which is freely available to use and adapt without restriction. Using the sequencing data from 3 human cell lines and 14 post-mortem brain tissue samples, we showed that our pipeline produces state-of-the-art SNP, structural variant and methylation calls at a cost comparable to short-read sequencing. This makes large-scale long-read sequencing projects feasible; the protocol is currently being used to sequence thousands of brain genomes as a part of the NIH CARD initiative.

We developed a new Hapdup method that generates *de novo* diploid assemblies from ONT sequencing only. We showed that outside of centromeres and segmental duplications, our assemblies are structurally highly concordant with the HPRC *de novo* assemblies that were produced from the more expensive combination of multiple sequencing technologies. Our assembly-based structural variant calls were highly accurate and complete, showing improvement over the current short-read and long-read methods. The assembly representation allowed us to construct pangenomes of the MHC and IGH regions that are highly polymorphic and are otherwise difficult to analyze with linear references. Multiple new approaches have been recently developed to to genotype SVs within larger collections of short-read samples using similarly constructed pangenomes, thus providing means to ascertain their frequencies in larger populations (Ebler et al., 2022; Sirén et al., 2021).

Our SNP calls produced with PEPPER-Margin-DeepVariant were comparable to or better than short-read-based methods. As expected, the most noticeable improvement was associated with the regions of low short-read mappability. Consistent with previous studies, we showed that using T2T-CHM13 as a reference genome increased the number of small variants that could be profiled. We also added a new functionality to Margin that can phase small and structural variant calls into megabase-scale haplotypes and reduces phasing switch error. Our methylation calls were highly concordant with the standard bisulfite sequencing, but in addition had haplotype-specific resolution, highlighted by our analysis of differentially methylated promoters.

Although an improvement over the previous ONT benchmarks, small indels inside homopolymers and low-complexity repeats remain the major source of the residual errors. This constitutes approximately two-thirds of all small indels in a human genome, but a minor fraction of overall variation in protein-coding sequence. Our evaluation of the R10 sequencing protocol showed substantial improvements in indel accuracy compared to the R9 protocol, in particular inside protein-coding regions. However, residual errors in long homopolymer and tandem repeat regions remain a challenge. In addition, ONT basecalling software remains one of the computational bottlenecks.

Our *de novo* diploid assemblies had high structural quality through most of the human genome, but did not reconstruct many long segmental duplications. This is primarily due to the reduced read length, compared to standard ONT protocols (most unassembled segmental duplications are longer than our read’s N50). However, it may be possible to reconstruct some of the missing duplications using a haplotype clustering approach (Vollger et al., 2019). In the current version of the pipeline, we called structural variants using *de novo* assemblies, but small variants were called using reference mapping. Further optimizations of ONT read error rate and improvements in our genotyping algorithms will allow for a more elegant solution to call phased small variants directly from diploid assemblies. We are also planning to integrate assemblies and methylation data better, which will allow methylation profiling of long sample-specific insertions.

Our analysis of structural variation calls highlighted that different methods may represent the same genomic variation differently, for example, by splitting or merging multiple indels in close proximity, or shifting alignment coordinates inside VNTRs. This results in difficulties in variant comparison across multiple samples or methods (English et al., 2022). *De novo* assemblies implicitly encode the genomic variation, and new pangenome graph methods (Liao et al., 2022) aim to provide an alternative representation of small and structural variations. However, reference mapping is still required for matching the variant calls against the existing databases (Chowdhury et al., 2022; Kirsche et al., 2021). Further improvements in structural variant representation and comparison models will be critical for the next large-scale, long-read genomic studies.

## Methods

### Sample collection

For the NABEC brain samples, frozen tissue was sampled from the frontal cortex for 14 neurologically normal individuals. All samples were obtained from the Banner Sun Health Research Institute (https://www.bannerhealth.com/services/research/locations/sun-health-institute/programs/body-donation/tissue). All individuals were of European ancestry and had no clinical history of neurological or cerebrovascular disease, or a diagnosis of cognitive impairment during life. Demographics, tissue source and cause of death for each subject are shown in Supplementary Table 1. Average age at time of death was 85.2 years of age (range, 68 - 95 years) and 8 were male and 6 were female.

For the cell lines, high molecular weight (HMW) DNA was extracted from the following cultured cell lines purchased from Coriell (https://www.coriell.org/): HG002 (Ashkenazi Jewish ancestry, cat. no. GM24385), HG02723 (African ancestry, cat. no. HG02723) and HG00733 (American ancestry, cat. no. HG00733). For these three cell-lines, cell culture was performed using Epstein–Barr virus (EBV)-transformed B lymphocyte culture from the cell-lines in RPMI-1640 media with 2 mM L-glutamine and 15 % fetal bovine serum at 37 °C.

### DNA processing

The frontal cortex (J Billingsley et al., 2022) and cell-lines (J Billingsley, 2022) protocols are explained in detail and are both publicly available on protocols.io. In brief, for the frontal cortex samples ∼40mg of frozen tissue was homogenized with a Tissueruptor instrument (Qiagen). HMW DNA was then extracted using a Kingfisher Apex instrument (Thermofisher) with a custom script and Nanobind Tissue Big DNA kit, which uses 3mm Nanobind disks (Circulomics/ PacBio, US). The HMW DNA was sheared to a target size of 30kb with the DNA Fluid + needles at speed 45 for two cycles on a Megaruptor3 instrument (Diagenode).

The cell-line HMW DNA was extracted manually from 4×10^6^ of cells using a Nanobind Tissue Big DNA kit (Circulomics/PacBio, US). An enrichment of short DNA fragments was observed for the Coriell cell-lines post DNA extraction. Therefore, to yield data comparable to the brain sequencing, using the Short Read Eliminator kit (SS-100-101-01) from Circulomics/Pacbio, a size selection step was included to deplete DNA fragments up to 25kb. Finally, the HMW DNA was then sheared to a target size of 30kb with the DNA Fluid + needles at speed 45 for two cycles (Diagenode). For all samples, DNA length was assessed by running 1ul on a genomic screentape on the TapeStation 4200 (Agilent). DNA concentration was assessed using the dsDNA BR assay on a Qubit fluorometer (Thermo Fisher).

### Library preparation

Libraries were constructed using an SQK-LSK 110 kit (ONT). To ensure minimal DNA loss during library preparation, and to retain long DNA fragments, the following modifications were made to the standard SQK-LSK 110 (ONT) protocol; 1) 4.5 ug of DNA was used as starting input. This is higher than the recommended amount due to the fact that around 60-75% of the starting material is lost during library preparation. Therefore, starting with 4.5ug DNA input ensured we could do three loads of 400ng with the final library, 2) to reach the 4.5ug DNA input the starting DNA volume was usually higher than the recommended 47ul. In this case, the volume of AMPure XP beads was modified to match the input DNA volume, i.e. if 58ul of DNA was added then 70ul was added of AMPure XP beads, 3) during DNA repair and end-prep, 75% ethanol was used for washing rather than 70% (per ONT, anything between 70-80% is acceptable), 4) at step 16 of DNA repair and end-prep, the original elution time was 2 minutes at room temperature. This was modified to 3 minutes at 37°C with light shaking on a Thermomixer instrument (Eppendorf) at 450rpm, 5) during Adapter ligation and clean-up, 45uL of AMPure XP beads were used 6) SFB was used, and finally 7) in step 16 of the Adapter ligation and clean-up, the final elution conditions were changed to 20 minutes at 37°C.

### PromethION sequencing

PromethION sequencing was performed as per manufacturer’s guidelines (ONT, FLO-PRO002) with minor adjustments, such as 400ng of the library was loaded onto each primed R9.4.1 flow cell to maximize data output. Over the three-day run, most samples only required one additional load (usually around 48 hours). However, some runs hit a pore occupancy of ∼2000 earlier, which was usually due to the variability across PromethION flow cells. In these cases, two reloads were required, which usually meant one reload every 24 hours.

### PEPPE-Margin-DeepVariant updates

We improved the variant calling speed and accuracy of PEPPER-Margin-DeepVariant by adding features that allow skipping the re-genotyping of candidate variants that are passed to DeepVariant. PEPPER first finds candidate variants and then genotypes them with a smaller recurrent neural network. The variants that are above a certain confidence threshold can be skipped from re-genotyping with a larger convolutional neural network used in DeepVariant. The detailed description of the updated methods is provided at the repository (https://github.com/kishwarshafin/pepper/blob/r0.8/docs/misc/pepper_methods.md). We used PEPPER-Margin-DeepVarian package r0.8 with default parameters for ONT reads.

### Small variant benchmarking

We used the Genome in a Bottle (GIAB) small variant benchmark to evaluate the performance on the HG002 genome, using benchmark v4.2.1 inside the confident intervals defined in “HG002_GRCh38_1_22_v4.2.1_benchmark_ noinconsistent.bed”. SNP recall and precision of ONT and Illumina variant calls were computed using hap.py v0.3.14 (https://github.com/Illumina/hap.py). Since the GIAB benchmark is only available for HG002, we also used variant calls produced with DeepVariant using HiFi reads as ground truth for HG002, HG00733, and HG02723. On HG002, both GIAB and DeepVariant call sets resulted in very similar statistics (Supplementary Table 2). We used the confident regions defined for HG002 for the other cell lines, which explains the slight decreases in recall and precision for both ONT and Illumina for HG00733 and HG02723.

To benchmark phase switch and hamming error rates, we used small variant calls produced from HPRC assemblies using dipcall v0.3 as ground truth. Error rates were then computed using “whatshap compare” v1.5 (Martin et al., 2016) module.

### Hapdup pipeline and assembly-based structural variant calling

Hapdup takes as input (i) a set of haploid contigs produced by any long-read assembler, such as Shasta or Flye and (ii) alignment of the original long-reads against the assembly produced by minimap2 (Li, 2018). Such assembly only contains ∼50% of the heterozygous variants, and our goal is to convert it into a locally phased diploid assembly that contains the complete set of structural variants in the genome.

Because de novo assemblies may leave some repetitive parts of the genome unassembled, reads originating from these unassembled repeat copies may misalign to their paralogs (if they happen to be assembled). Because the copies of long repeats often are not exact, misaligned reads may create artificial “haplotypes”, in addition to the (correctly mapped) paternal and maternal alleles. It is important to filter out misaligned reads to ensure that the subsequent diploid phasing is correct. Hapdup filters out reads with either (i) large unaligned parts or (ii) high alignment error. Afterwards, PEPPER is used to call SNPs, SNPs are phased using Margin, and reads are haplotagged according to their phases.

Next, Hapdup runs two instances of the Flye polisher, using only the aligned reads from either first or second phase. The polisher algorithm separates the input alignment into small chunks, and each chunk is polished reference-free using the maximum likelihood approach (Y. Lin et al., 2016). To recover a large heterozygous variant, it is important that reads containing the variant are consistently aligned, which may be difficult in VNTR regions. To ensure the correct splitting of the original alignment, we added new functionality to the polishing algorithm that identifies regions with indels with inconsistent coordinates between reads, and ensures that the problematic regions are contained inside a single mini-alignment.

The polishing procedure can recover indels and substitution variants, but was not designed to detect structural variants that are created by genomic rearrangements (for example, inversions). Instead, Hapdup detects genomic rearrangement signatures from the read alignments, infers their phases, and applies the rearrangement to the corresponding set of contigs.

Hapdup outputs phased haplotypes in two different formats. First, it generates a “dual” assembly, that has the same contiguity as the original haploid set of contigs, but may contain occasional phase switches if phase blocks are shorter than contigs. Alternatively, Hapdup can split the contigs at the regions that lack proper phasing (indicated by Margin), so that every contig represents a contiguous paternal or maternal haplotype.

We used Shasta v0.10.0 with config “Nanopore-CARD-Jan2022.conf” and Hapdup v0.11 assemblies. Assemblies were also evaluated using QUAST v5.2.0 (Mikheenko et al., 2018), yak (https://github.com/lh3/yak), asmgene (a part of paftools v2.24-r1152-dirty; (Cheng et al., 2021, 2022). We called SNPs from assemblies using dipcall v0.3 and confirmed a low switch error rate of 0.07-0.18% using the “whatshap compare” v1.5 (Martin et al., 2016) module.

### Misassembly detection using Flagger

To detect misassemblies in the Shasta+Hapdup and HPRC assemblies, we ran Flagger v0.2.0 (Liao et al., 2022). In summary, Flagger takes the read alignment to a diploid/dual assembly, computes read depth of coverage, fits a Gaussian Mixture Model to the coverage distribution and identifies the best coverage thresholds for partitioning the assembly into 4 main components; erroneous, falsely duplicated, haploid (correctly assembled) and collapsed. To compute read depth of coverage along the assemblies, we aligned both HiFi and ONT reads to each LCL assembly including both haplotypes as the reference. We used the alignment WDL available in the Flagger repository on Dockstore (https://dockstore.org/my-workflows/github.com/mobinasri/flagger/LongReadAlignerScattered). The workflow parameters were set as follows: *preset = “map-pb” / “map-ont”; aligner = “winnowmap”; options = “--cs --eqx -Y -L -I8g -p0.1”; kmerSize = 15*.

To fix probable phasing issues in the read alignments, we applied Secphase (v0.2.0) on both ONT and HiFi alignments before running Flagger. Secphase calculates a marker consistency score for each alignment and selects the top alignment among all primary and secondary alignments of a single read. If the selected alignment is secondary, the primary tag will be assigned to this alignment. The marker consistency score is based on a multiple sequence alignment of all the blocks one read is aligned to (Liao et al., 2022).

While running Winnowmap, we changed the parameter -p from 0.8 (default value) to 0.1 to include more secondary alignments per read. Therefore, Secphase will potentially have additional alignment candidates to select from. A new parameter, *--primMarginRandom*, was added to Secphase in the latest version 0.2.0. This parameter allows Secphase to randomly select one alignment if the difference between scores of the original primary alignment and the highest secondary alignment is lower than this given threshold. This parameter was useful in retrieving alignments in highly homozygous regions. This change was made because we observed that one haplotype would have all of the reads mapped to it while the other haplotype would be almost free of any alignment due to a small number of errors in the unmapped haplotype; however, after running Secphase with this change those alignments became equally distributed between the two haplotypes. For HiFi data we used ‘--hifi’ preset, which has *--primMarginRandom* 50 as default and for ONT we used ‘--ont’ preset with *--primMarginRandom* 10 as default. Supplementary Figure 14 shows an example of how changing -p and using Secphase could improve the alignment to the diploid assembly.

For running Secphase, we used the workflow available in the Secphase repository on Dockstore. (https://dockstore.org/my-workflows/github.com/mobinasri/secphase/Secphase). The workflow parameters were set as follows: *secphaseOptions = “--ont” / “--hifi”; secphaseDockerImage = “mobinasri/secphase:v0.2.0”.* The output of this workflow will be a log file, outLog, containing the name of the reads whose alignments have to be relocalized. This log file will be passed to the next workflow FlaggerPreprocess.

Prior to running Flagger, we ran the FlaggerPreprocess workflow, which is used for applying the alignment modifications suggested by Secphase, removing highly divergent alignments, calling variants (presumably caused by assembly errors or mismapping), removing alignments with alternative alleles in those variants, and calculating read coverage along the assembly both before and after removing alternative-contained alignments. To give a summary of alignment removal, FlaggerPreprocess has a step that extracts high-quality SNPs, iterates over each alignment and if an alignment contains the alternative allele of those SNPs it will be removed since this is frequently the result of mismapping. If assemblies are not assembled/polished with the reads of high base-level accuracy like HiFi or Illumina reads, this step will make a lot of regions free of any alignment. The workflow provides two coverage files, one including all of the alignments and the other one after excluding alternative-contained alignments. Our main goal here is using the coverage file that is derived from all of the alignments. We also used the filtered version of the HiFi alignments.

For running the preprocessing steps, we used the workflow available in the Flagger repository (https://dockstore.org/my-workflows/github.com/mobinasri/flagger/FlaggerPreprocess) The workflow parameters were set as following: *minReadLength = 5000; minAlignmentLength = 5000;* maxDivergence = 0.02; variantCaller = “dv” for HiFi; maxDivergence = 0.09, variantCaller = “pmdv” for ONT.

Flagger also needs a list of bed files pointing to the regions that are prone to have coverage biases. ONT and HiFi coverage biases in some Human Satellites. Based on those observations, Flagger repository provides a list of bed files for HiFi and ONT. Two bed files are related to the regions which may have increasing (mostly HSat2/3) and decreasing (mostly HSat1) biases in HiFi coverage, and one bed file is for decreasing (mostly HSat1) bias in ONT coverage. These bed files in the CHM13 coordinates are available in the Flagger github repository (https://github.com/mobinasri/flagger/tree/main/misc/biased_regions). Coverage biases may not exist in all datasets but Flagger increases the degree of freedom of its model to avoid misflagging if such biases exist.

To project the blocks from CHM13 coordinates to the assembly coordinates, we used another workflow provided by Flagger (https://dockstore.org/my-workflows/github.com/mobinasri/flagger/ProjectBlocksForFlagger). This workflow takes a list of bed files along with the alignment of each haplotype assembly to the CHM13 reference and outputs the projections of those bed files to each haplotype. This workflow also takes three other bed files for sex chromosomes, centromeric regions and segmental duplications. Each one of them will be projected similarly and their projection will be used in running the FlaggerStats workflow to stratify the Flagger results by these annotations.

For running the Flagger core pipeline, we used the workflow available in the Flagger repository on Dockstore (https://dockstore.org/my-workflows/github.com/mobinasri/flagger/Flagger). The coverage files and the projection of the biased bed files produced by FlaggerPreprocess and ProjectBlocksForFlagger will be the inputs to the Flagger workflow. It will output a bed file that partitions each diploid assembly into 4 main components; erroneous, falsely duplicated, haploid (correctly assembled) and collapsed. Any component other than haploid indicates a type of misassembly.

The workflow parameters were set as follows. For HiFi: biasedRegionBedArray = [${HiFi_High_Coverage_BED_PAT}, ${HiFi_High_Coverage_BED_MAT}, ${HiFi_Low_Coverage_BED_PAT}, ${HiFi_Low_Coverage_BED_MAT}]; biasedRegionNameArray = [“hifi_high_pat”, “hifi_high_mat”, “hifi_low_pat”, “hifi_low_mat”]; biasedRegionFactorArray = [1.25, 1.25, 0.75, 0.75]. For ONT: biasedRegionBedArray = [${ONT_Low_Coverage_BED_PAT}, ${ONT_Low_Coverage_BED_MAT}]; biasedRegionNameArray = [“ont_low_pat”, “ont_low_mat”]; biasedRegionFactorArray = [0.75, 0.75];

The output bed file of Flagger with the projected annotations (including sex, centromere and SD) was then given to the FlaggerStats workflow to produce tables of statistics (https://dockstore.org/my-workflows/github.com/mobinasri/flagger/FlaggerStats).

### Structural variant calling and benchmarking

To produce structural variant calls from diploid assemblies, we developed a tool called hapdiff, which is based on a modified version of SVIM-asm (Heller & Vingron, 2020). The package incorporates minimap2 with predefined alignment parameters, which were optimized for alignment of regions containing long structural variants (“-ax asm20 -B 2 -E 3,1 -O 6,100”). Having a fixed set of alignment parameters is also critical for reproducibility. Second, we added the functionality to group the variants inside the same VNTR together, if the annotation is provided. As illustrated in Supplementary Figure 5, VNTR grouping substantially improves the agreement between Hapdup and HPRC assemblies. This highlights the challenge of ensuring alignment consistency inside VNTR regions.

Sniffles2 was run using default parameters, with a VNTR annotation file provided. CuteSV was run with parameters recommended for ONT datasets (*--max_cluster_bias_INS 100 --diff_ratio_merging_INS 0.3 --max_cluster_bias_DEL 100 --diff_ratio_merging_DEL 0.3*)

We used hapdiff v.0.7. Multiple sets of structural variants were compared using Truvari v3.3.0 with added “-r 2000” parameter that controls the maximum linear distance between two variants. We also tested Truvari with default parameters and found that it has only a small effect on the resulting F1 scores and affected different tools similarly (Supplementary Table 4).

### Harmonized phased representation of small and structural variants

Modifications to Margin were made to support joint phasing of small and structural variants. The underlying phasing algorithm remained the same: allele support from reads is modeled by (i) selecting a configurable number of basepairs up- and downstream from the variant site from the reference, modifying this sequence to reflect each proposed allele, (ii) selecting a read subsequence based on the pairwise alignment to the extracted reference subsequence, and (iii) aligning each read subsequence to all alleles at each locus to generate emission probabilities for the HMM.

When experimenting with real data, we found that the proposed breakpoints for SVs would not always match the exact read alignment determined by the aligner. To account for this, we modified Margin to have two parameters for the distance up- and downstream from the variant for subsequence extraction: one for small variants (referenceExpansionForSmallVariants) and one for structural variants (referenceExpansionForStructuralVariants). A third parameter (indelSizeForSVHandling) was introduced to distinguish between small and structural variants with a configured value of 50bp. The small variant reference expansion value was left unchanged at 12bp. After experimentation on HG002’s chr20, we found that the reference expansion for structural variants had the greatest effect on accuracy at 64bp. Larger expansions did not yield notable phasing improvements but had increased runtime, while smaller expansions reduced phasing accuracy.

We used Margin (commit bb1e16a) with the config file “allParams.phase_vcf.ont.sv.json” to generate the final phased vcf files.

### Methylation calling

We produced phased 5-methylcytosine calls aligned to GRCh38 and T2T-CHM13 references. Initial methylation calls were produced *de novo* using Remora (https://github.com/nanoporetech/remora) incorporated in Guppy 6.1.2; afterward reads were aligned and phased using PEPPER-Margin-DeepVariant (Shafin et al., 2021) and annotated with modbamtools v0.4.8 (Razaghi et al., 2022) and modbam2bed v0.6.3 (https://github.com/epi2me-labs/modbam2bed). Regional haplotype methylation in gene promoters and SVs were calculated using “modbamtools calcMeth”.

### Pipeline implementation

The pipeline was implemented using the Workflow Description Language (WDL), and the corresponding tools are available as Docker containers. This implementation improves reproducibility and can automatically scale to different server or cloud environments. We ran our analysis using the Terra platform, running Cromwell (https://cromwell.readthedocs.io/) and using Google Cloud Platform as a backend. The pipeline could also be run on a local machine with a WDL server installed. We used the pipeline version f8ec6f2 (github commit).

## Code availability

The complete pipeline implemented in WDL is available at https://github.com/nanoporegenomics/card_nanopore_wf. Hapdup is available as a standalone tool at: https://github.com/KolmogorovLab/hapdup. Hapdiff is available at: https://github.com/KolmogorovLab/hapdiff.

## Data availability

The cell line data (HG002, HG0073 and HG02723) and openly available through the Terra workspace: https://anvil.terra.bio/#workspaces/anvil-datastorage/ANVIL_NIA_CARD_Coriell_Cell_Lines_Open. Human brain sequencing datasets are under controlled access and require a dbGap application (phs001300.v4). Afterwards, the data will be available through the restricted Terra workspace: https://anvil.terra.bio/#workspaces/anvil-datastorage/ANVIL_NIA_CARD_LR_WGS_NABEC_GRU

## Acknowledgements

This work was supported in part by the Intramural Research Program of the National Cancer Institute (NCI), the National Human Genome Research Institute (NHGRI), National Institute on Aging (NIA), and the Center for Alzheimer’s and Related Dementias (CARD), within the Intramural Research Program of the NIA and the National Institute of Neurological Disorders and Stroke (ZIANS003154, ZIAAG000538), National Institutes of Health (AG000538). This work utilized the computational resources of the NIH HPC Biowulf cluster (https://hpc.nih.gov). We thank members of the North American Brain Expression Consortium (NABEC) for providing samples derived from brain tissue. We are grateful to the Banner Sun Health Research Institute Brain and Body Donation Program of Sun City, Arizona for the provision of human biological materials. The Brain and Body Donation Program has been supported by the National Institute of Neurological Disorders and Stroke (U24 NS072026 National Brain and Tissue Resource for Parkinson’s Disease and Related Disorders), the National Institute on Aging (P30 AG19610 and P30AG072980, Arizona Alzheimer’s Disease Center), the Arizona Department of Health Services (contract 211002, Arizona Alzheimer’s Research Center), the Arizona Biomedical Research Commission (contracts 4001, 0011, 05-901 and 1001 to the Arizona Parkinson’s Disease Consortium) and the Michael J. Fox Foundation for Parkinson’s Research. B.P. was partly supported by NIH grants: R01HG010485, U24HG010262, U24HG011853, OT3HL142481, U01HG010961, and OT2OD033761. We acknowledge the support of Oxford Nanopore Technologies staff in generating this data set, in particular A. Markham. We also acknowledge the support of the Circulomics Inc team in generating this protocol, in particular K. Liu, J. Burke, M. Kim & D. Kilburn.

## Supplementary Tables

**(see attached Excel spreadsheets)**

**Supplementary Table 1.** Information about the ONT sequencing runs and computational pipeline performance.

**Supplementary Table 2.** Additional information about benchmarking small variant calls.

**Supplementary Table 3.** Assembly statistics and benchmarks.

**Supplementary table 4.** Structural variant benchmarks

**Supplementary table 5.** Methylation analysis statistics

**Supplementary table 6:** Analysis of rare SVs in brain samples

## Supplementary Figures

**Supplementary Figure 1.**
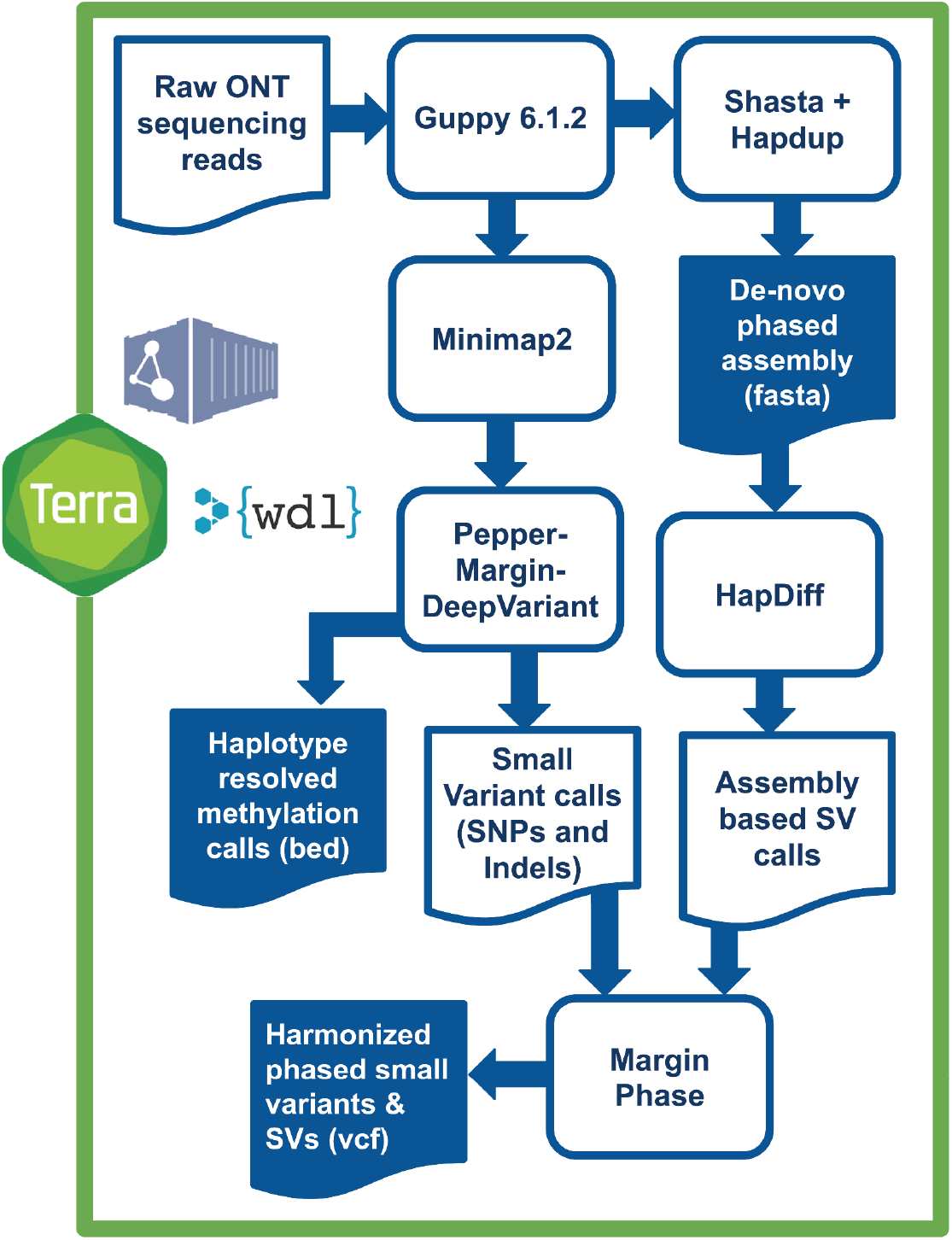
Overview of variant calling and methylation analysis pipeline. Raw ONT sequencing reads are basecalled by Guppy 6.1.2, which simultaneously produces methylation tags. A diploid, de-novo phased assembly is produced using a combination of Shasta and HapDup. These assemblies are used to call structural variants with HapDiff. Small variants are called against a reference genome with Pepper-Margin-DeepVariant. The phased alignment file generated by Margin is used to produce haplotype-resolved methylation calls. Small and structural variants are jointly phased by Margin, producing a single harmonized vcf.

**Supplementary Figure 2.**
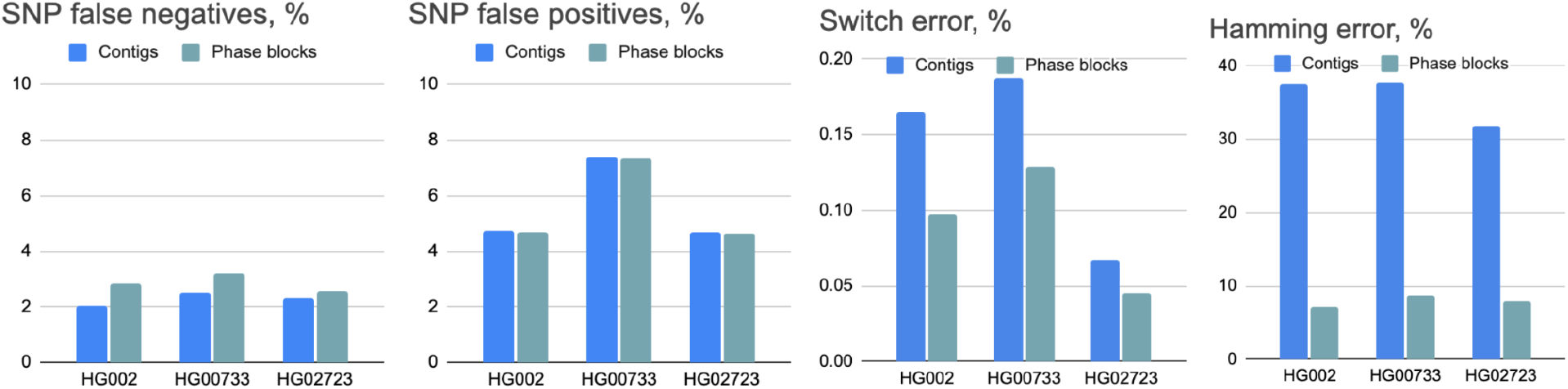
Assemblies switch and hamming errors computed using “whatshap compare”. SNPs were called using dipcall and benchmarked against small variant calls in HPRC assemblies.

**Supplementary Figure 3.**
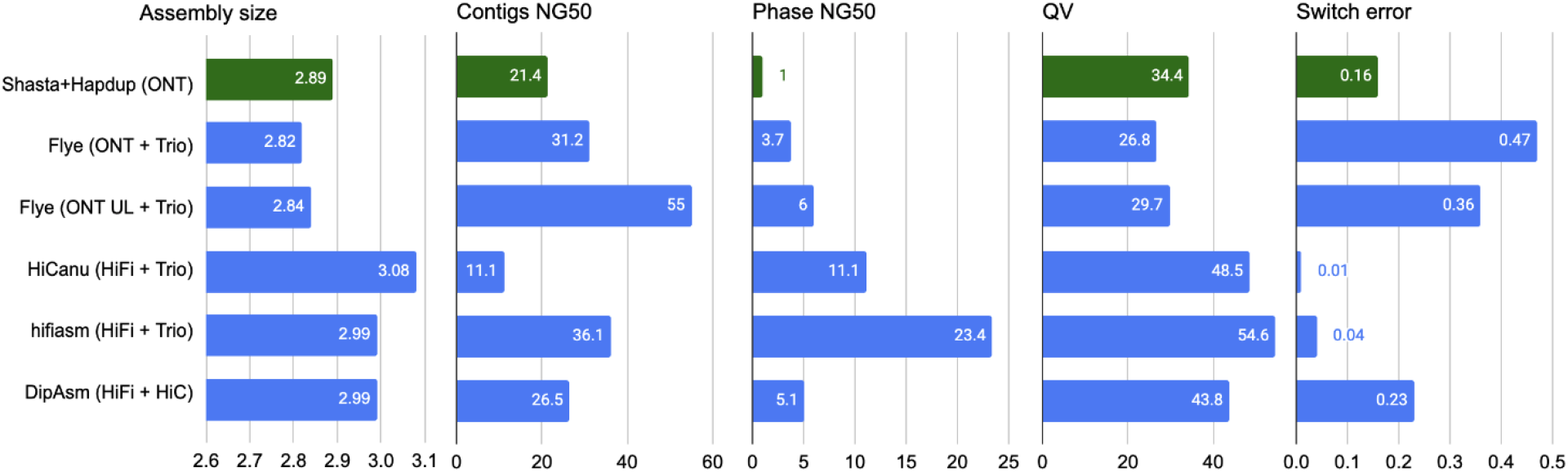
Assembly metrics comparison against HG002 assemblies produced in Jarvis et. al (2022). Our assemblies are highlighted in green. Flye (ONT+Trio) were produced using standard ONT reads at 60x coverage and Illumina parental information; Flye (ONT UL + Trio) is similar, but using ultra-long ONT extraction. HiCanu and hifiasm used 34x HiFi reads and Illumina parental sequencing. DipAsm used 34x HiFi reads and 60x Hi-C reads. Original evaluations from Jarvis et al. are shown. See Supplementary Table 3 for more detail.

**Supplementary Figure 4.**
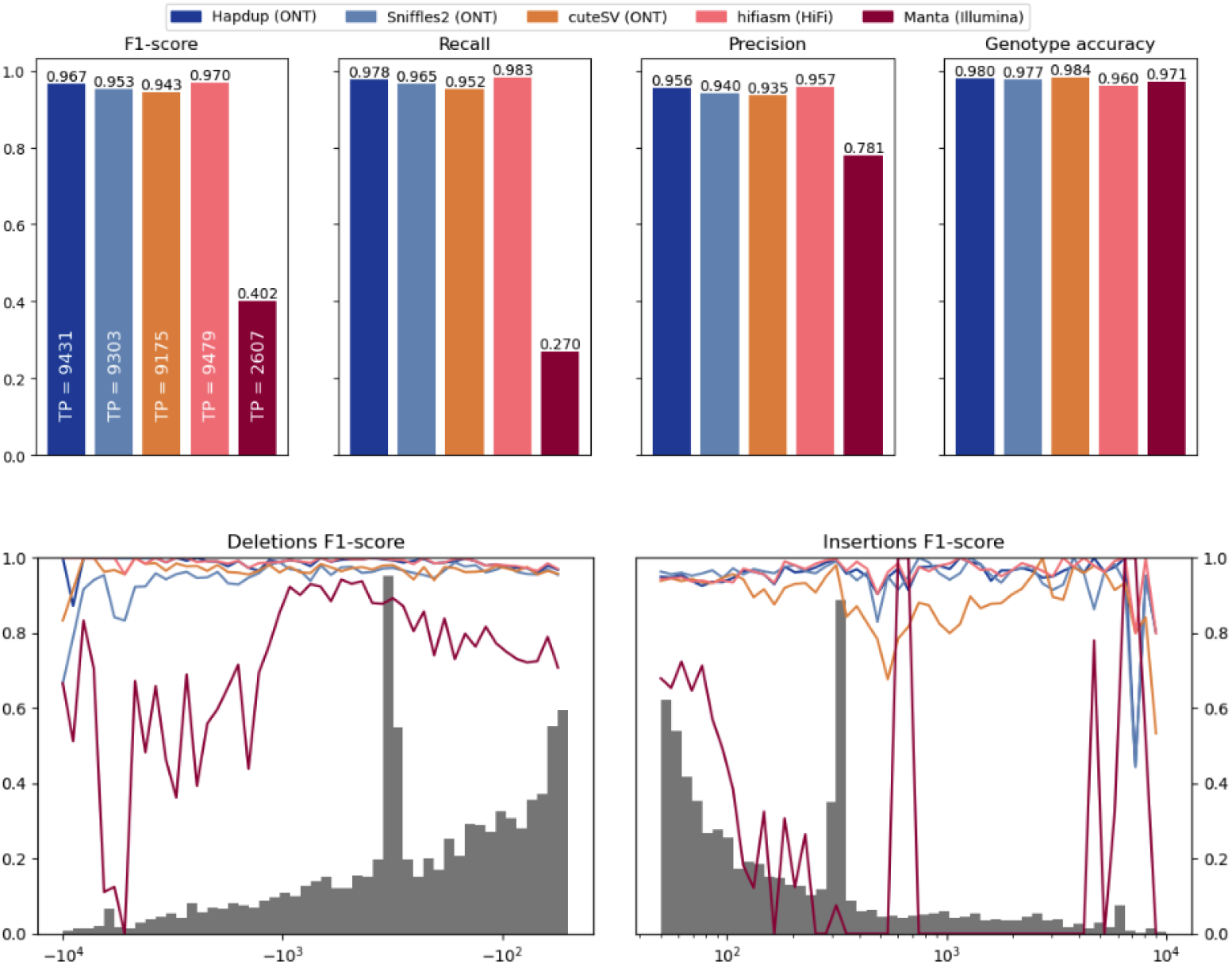
Comparison of structural variants using GIAB HG002 benchmark, including Illumina-based calls produced with Manta.

**Supplementary Figure 5.**
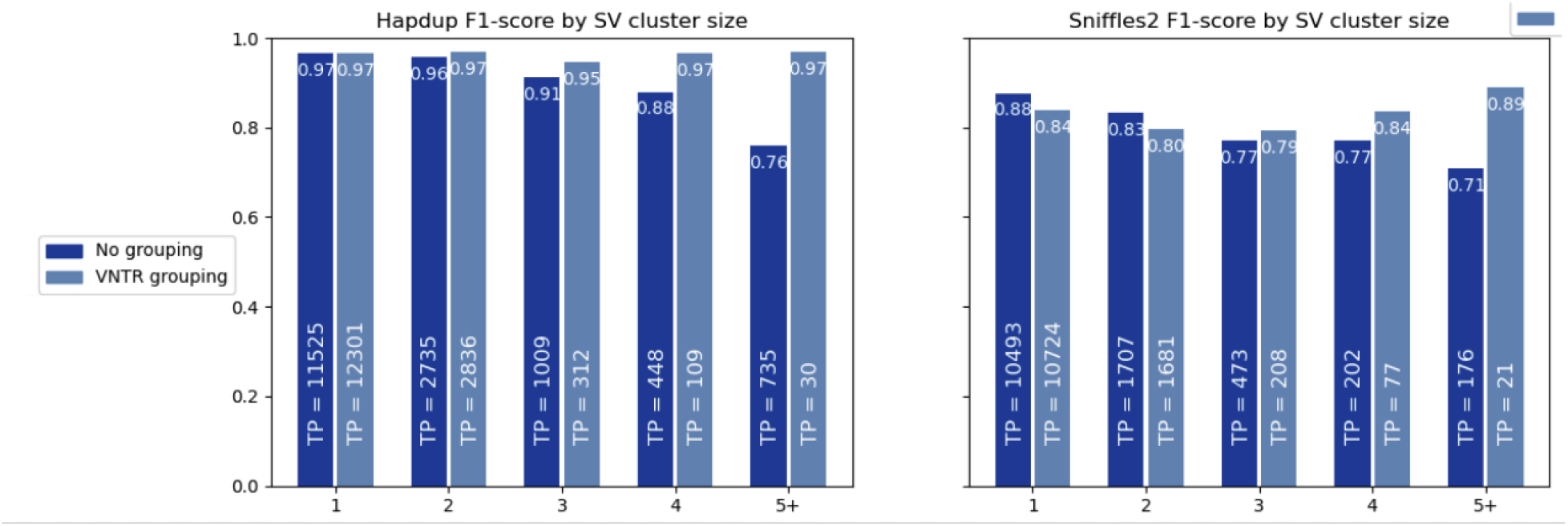
F1-score for SV inside clusters of different sizes. The HiFi calls for HG002 genome were used as reference, and calls within 2 kbp were clustered using single linkage clustering. The number of true positive calls in each category is shown as text. When VNTR grouping is enabled, all insertions and deletions within the same haplotype in a single VNTR are combined into a single call. A substantial portion of the reduced Sniffles2 concordance is explained by the differences in representation of SV clusters by the assembly-based and mapping-based approaches.

**Supplementary Figure 6.**
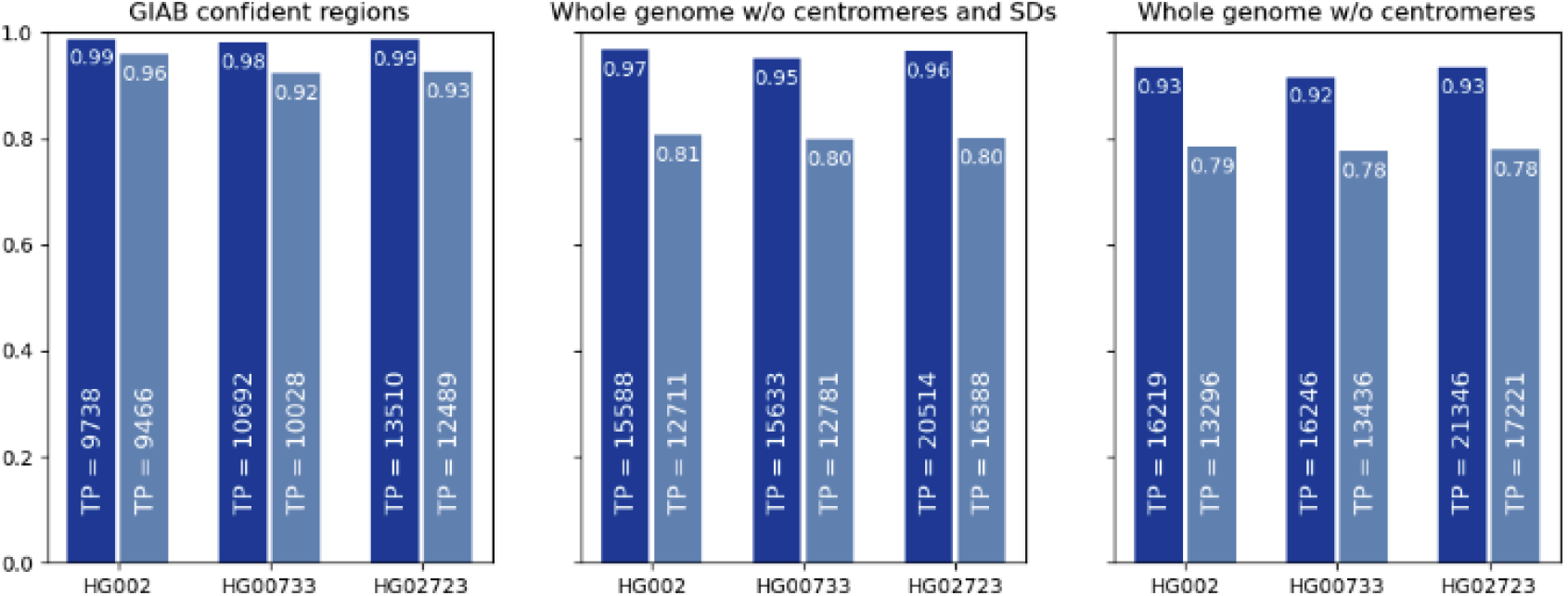
Comparison of Hapdup and Sniffles2 F1-scores against HiFi assemblies.

**Supplementary Figure 7.**
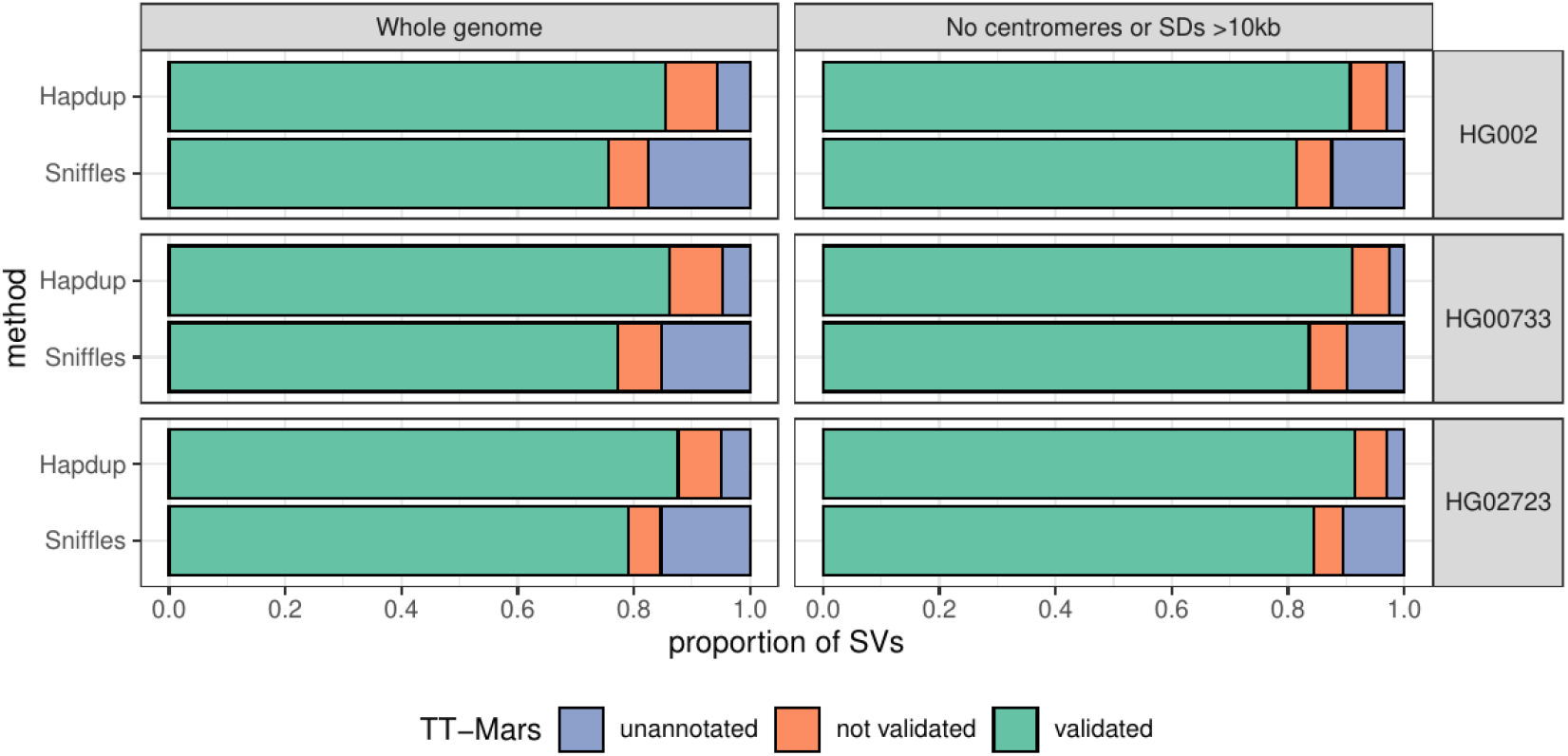
TT-Mars evaluation of Hapdup and Sniffles2 calls. Structural variant calls from Hapdup and Sniffles2 were compared to the assemblies from the HPRC for HG002 (top), HG00733 (middle), and HG02723 (bottom) with TT-Mars. The calls were either validated by the alignment (green), not validated (orange), or couldn’t be annotated by TT-Mars (blue). We evaluated all SVs across the genome (left), as well as the subset of SVs that don’t overlap centromeres or segmental duplications larger than 10 Kbp (right).

**Supplementary Figure 8.**
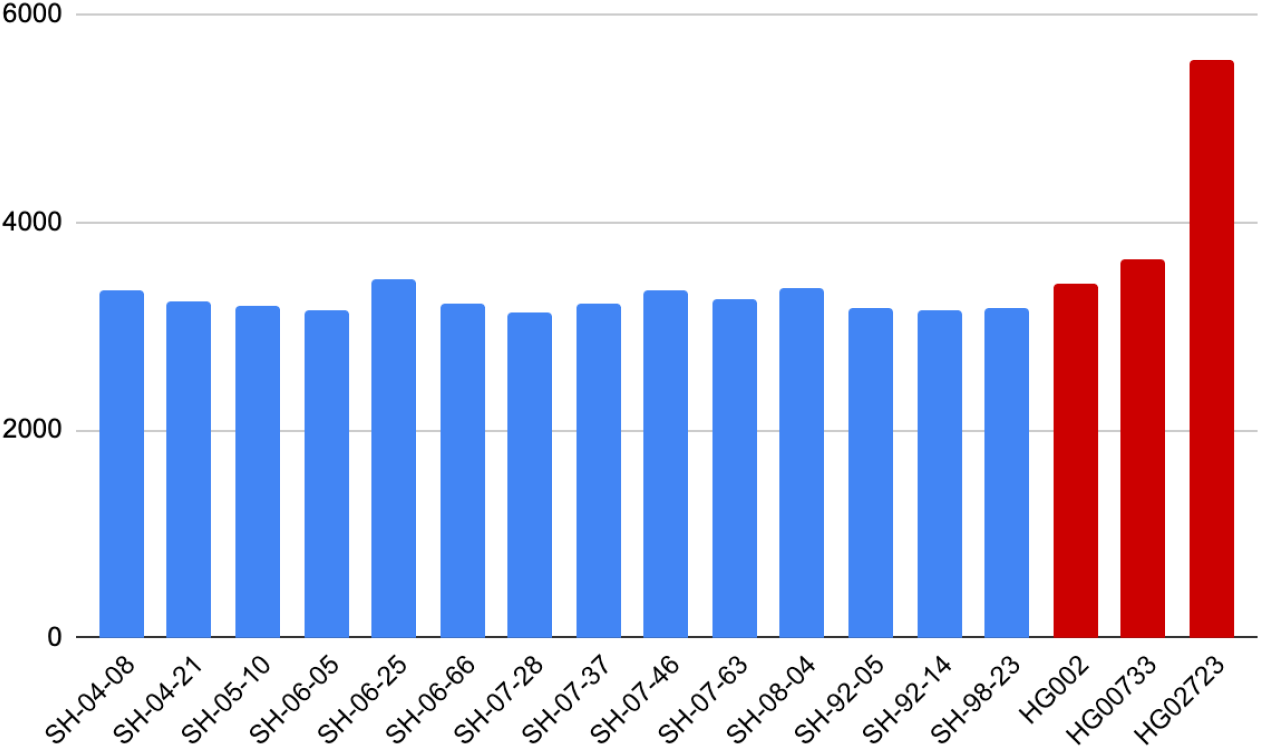
Number of SVs inside clusters of size at least 2. SVs are clustered with single linkage clustering within 2kb. Multiple indels within a single VNTR element are considered as single SVs in this analysis.

**Supplementary Figure 9.**
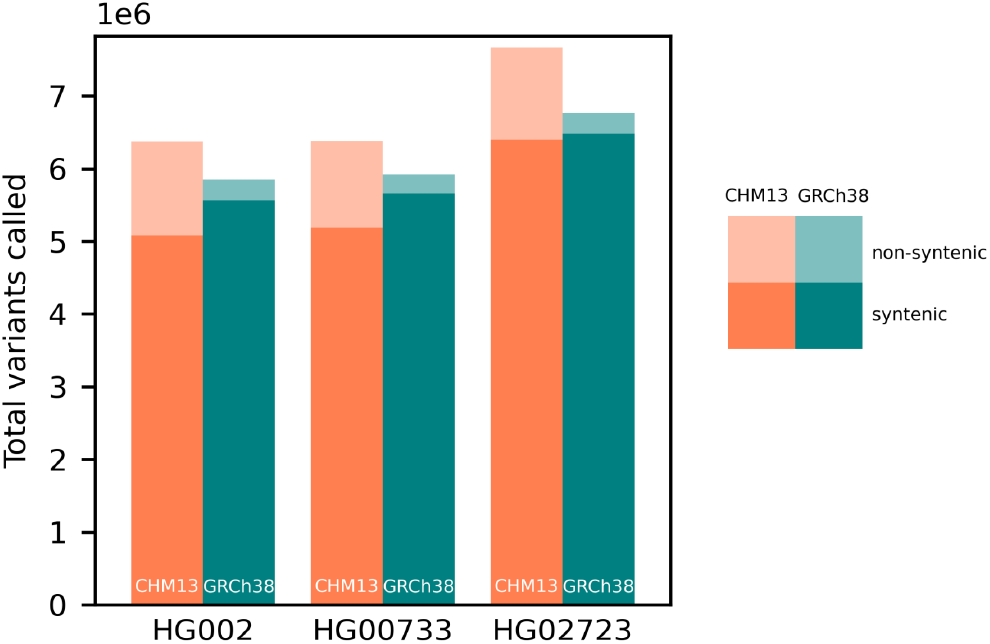
Comparison of small variant calling using GRCh38 and T2T-CHM13 references.

**Supplementary Figure 10.**
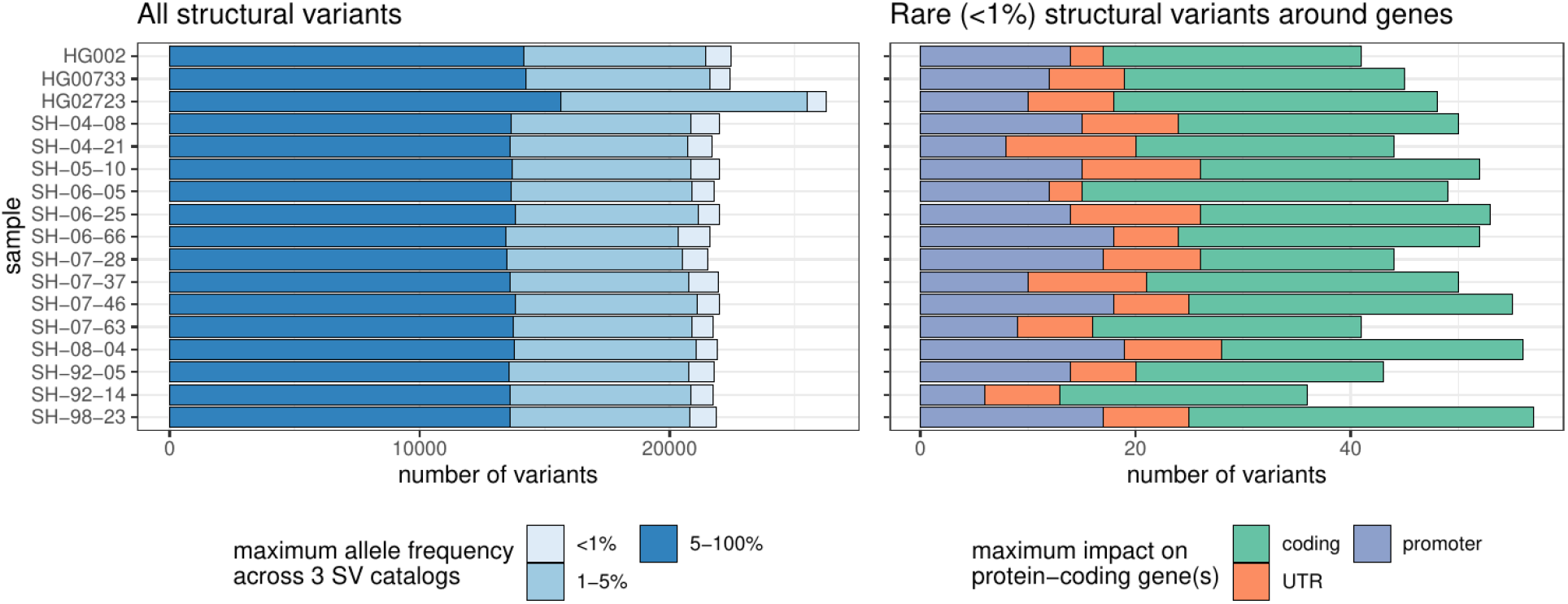
Lenient structural variant catalog. Similar to Figure 7A but including SVs close to centromeres, telomeres, or within segmental duplications were removed. (A) Number of structural variants across samples. In the left panel, structural variants were annotated with three SV catalogs (the gnomAD-SV database, a long-read-based SV catalog, and the HPRC v1.0 SV catalog). SVs are matched if they have at least 10% genomic overlap. The colors highlight the maximum frequency across these catalogs, the lighter blue showing “rare” SVs (with an allele frequency below 1%) in the catalogs, or unmatched. SVs may be unmatched, either because they are novel or due to the difficulties in the database comparison. The right panel shows the number of rare structural variants in protein-coding genes, grouped by their impact on the gene structure.

**Supplementary Figure 11.**
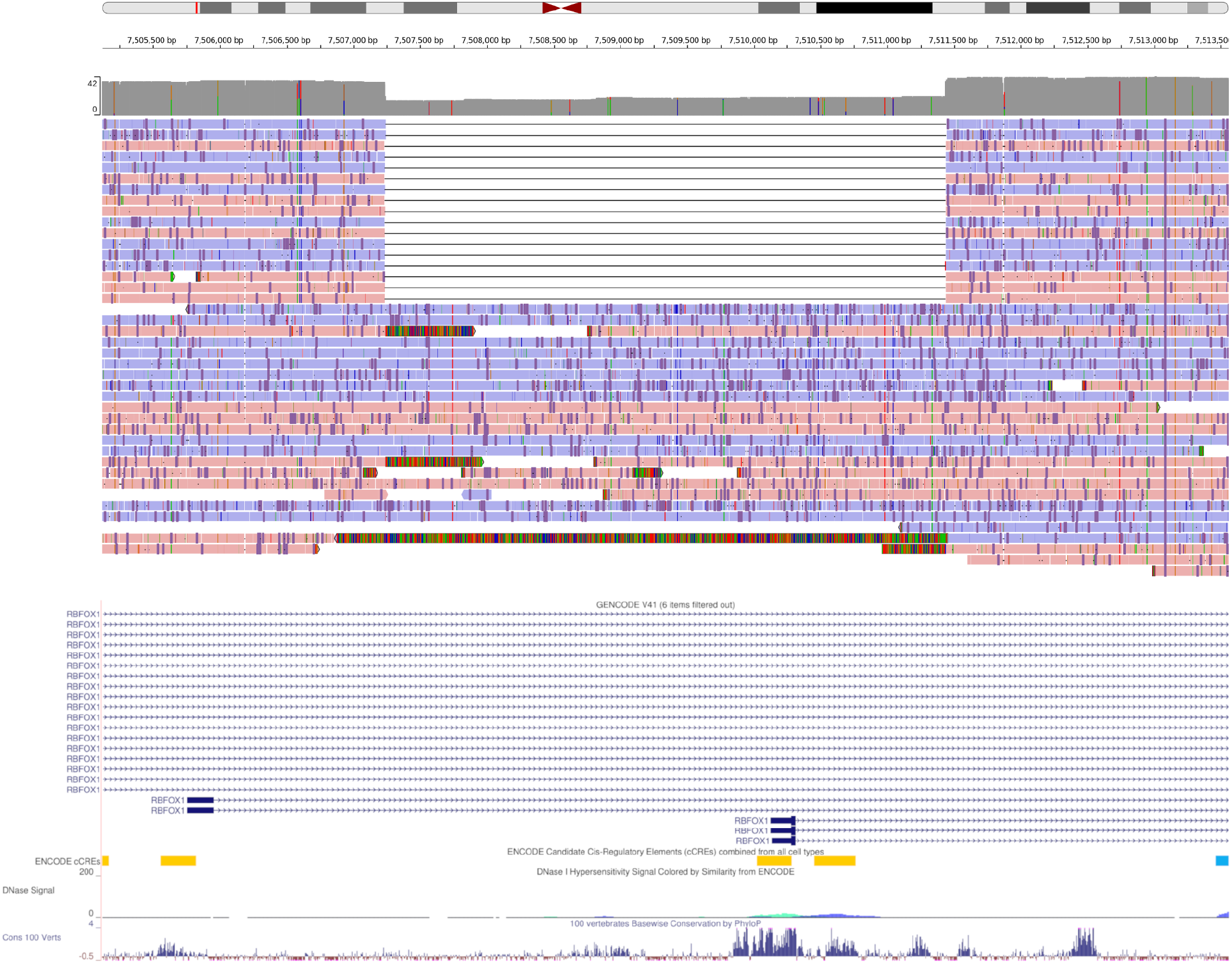
IGV view of a 4.2 Kbp heterozygous deletion of a transcription start site and exon of *RBFOX1*. The coverage histogram (dark grey) shows the drop in read coverage. The alignment of about half of the reads, labelled by strand (red/blue), support the deletion. The GENCODE track, ENCODE candidate cis-regulatory elements, and conservation tracks are shown at the bottom.

**Supplementary Figure 12.**
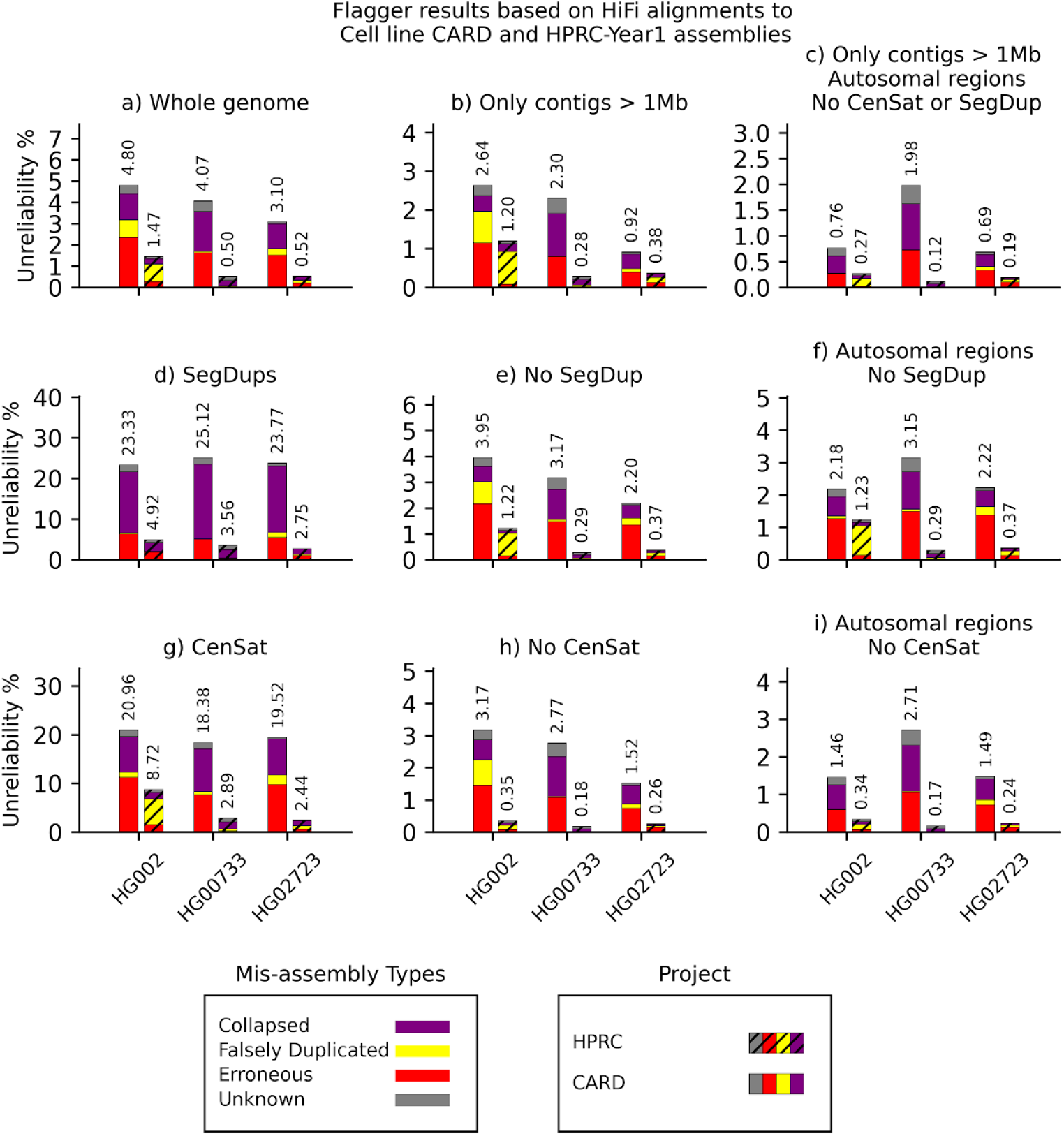
Flagger results based on HiFi alignments to cell line CARD and HPRC-Y1 assemblies. The y-axis of each panel indicates the unreliability percentages which are the total number of bases flagged as misassembly divided by the total assembly length and multiplied by one hundred

**Supplementary Figure 13.**
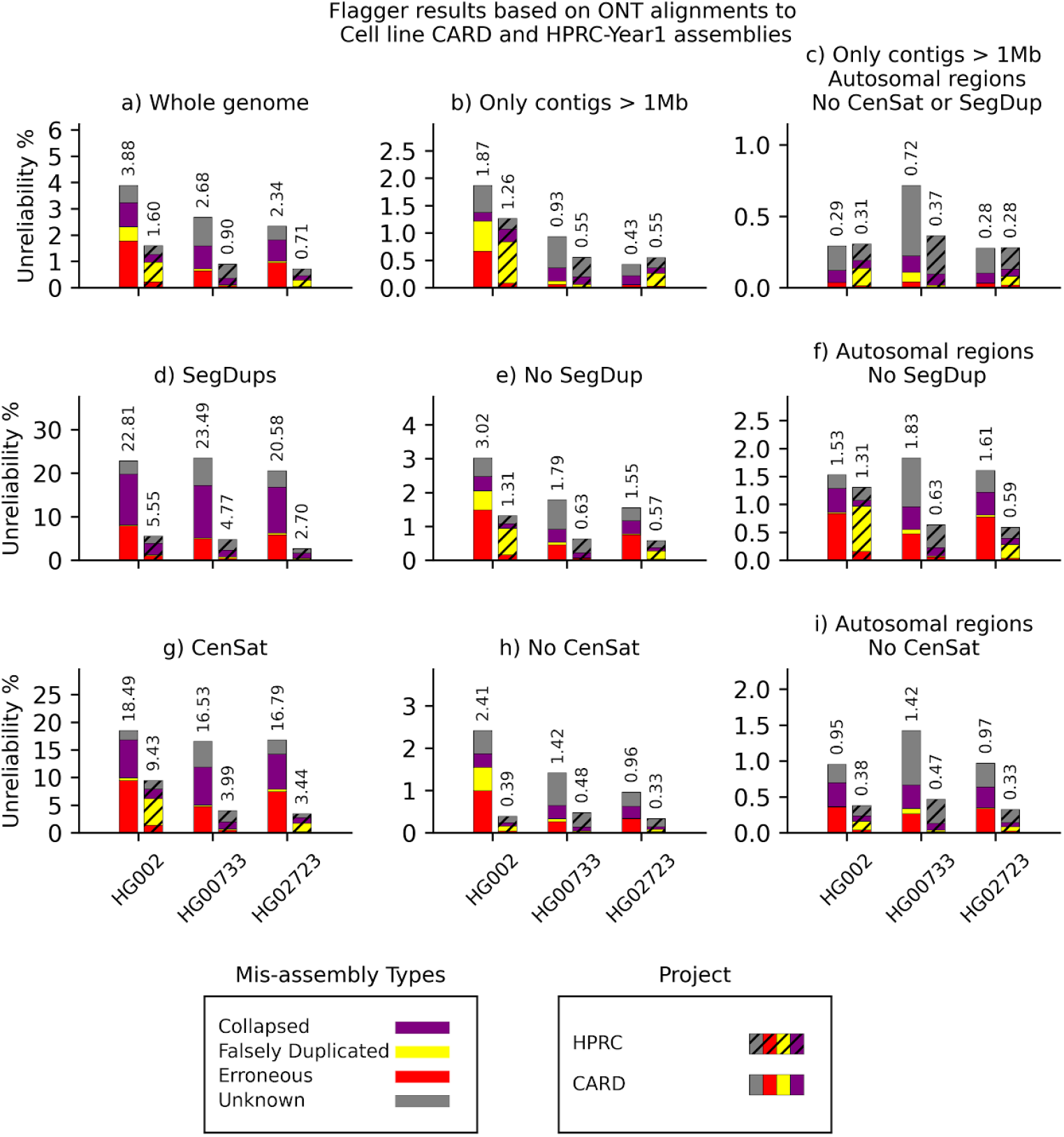
Flagger results based on ONT alignments to cell line CARD and HPRC-Y1 assemblies. The y-axis of each panel indicates the unreliability percentages which are the total number of bases flagged as misassembly divided by the total assembly length and multiplied by one hundred.

**Supplementary Figure 14.**
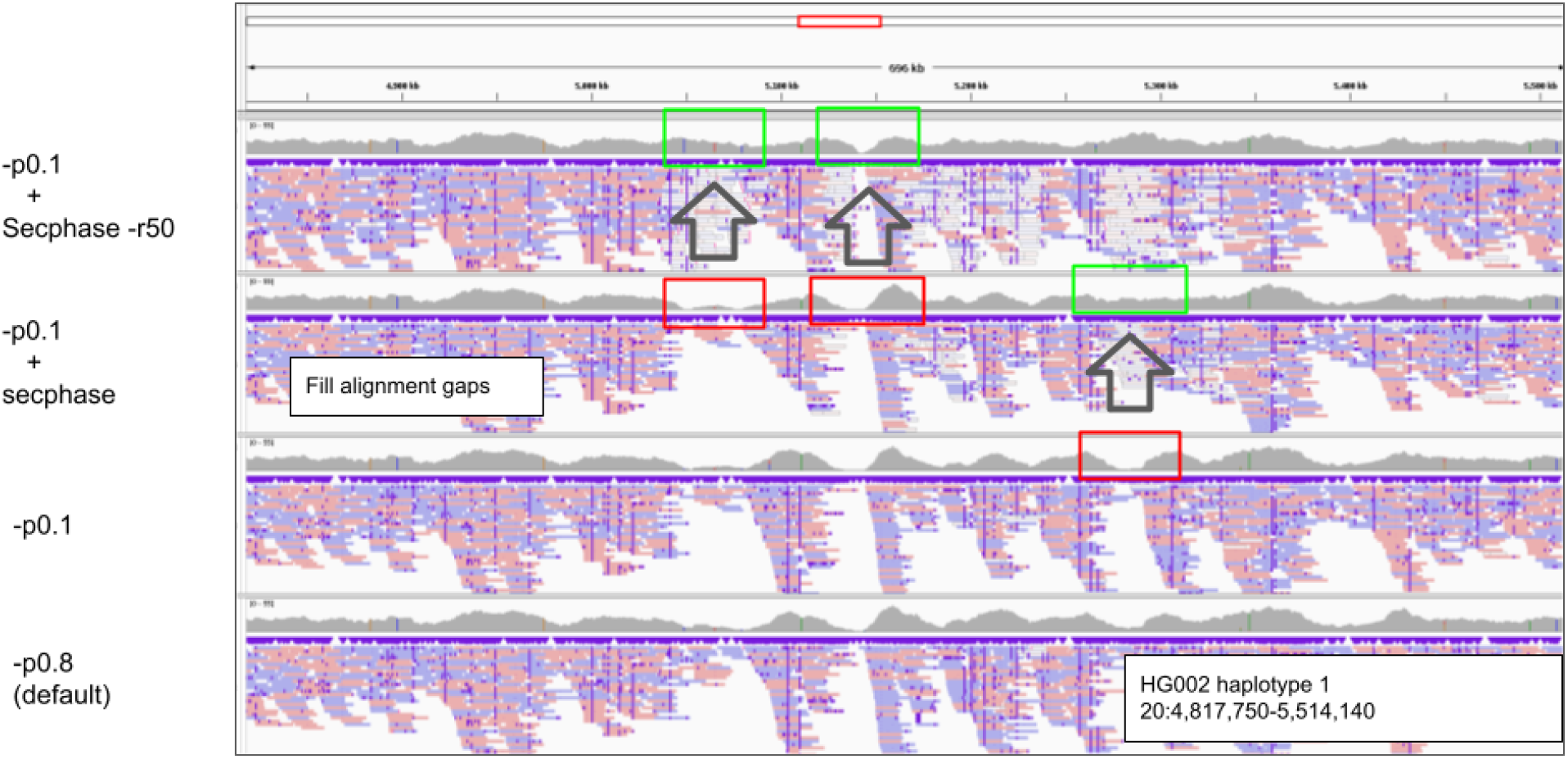
Adopting Flagger to ONT reads. Changing -p0.8 to -p0.1 increased the number of secondary alignments and filled some alignment gaps. Secphase with -r50 could not fill more of the alignment gaps with secondary alignments. The parameter -r50 was added to randomly assign reads in highly homozygous regions.

**Supplementary Figure 15.**
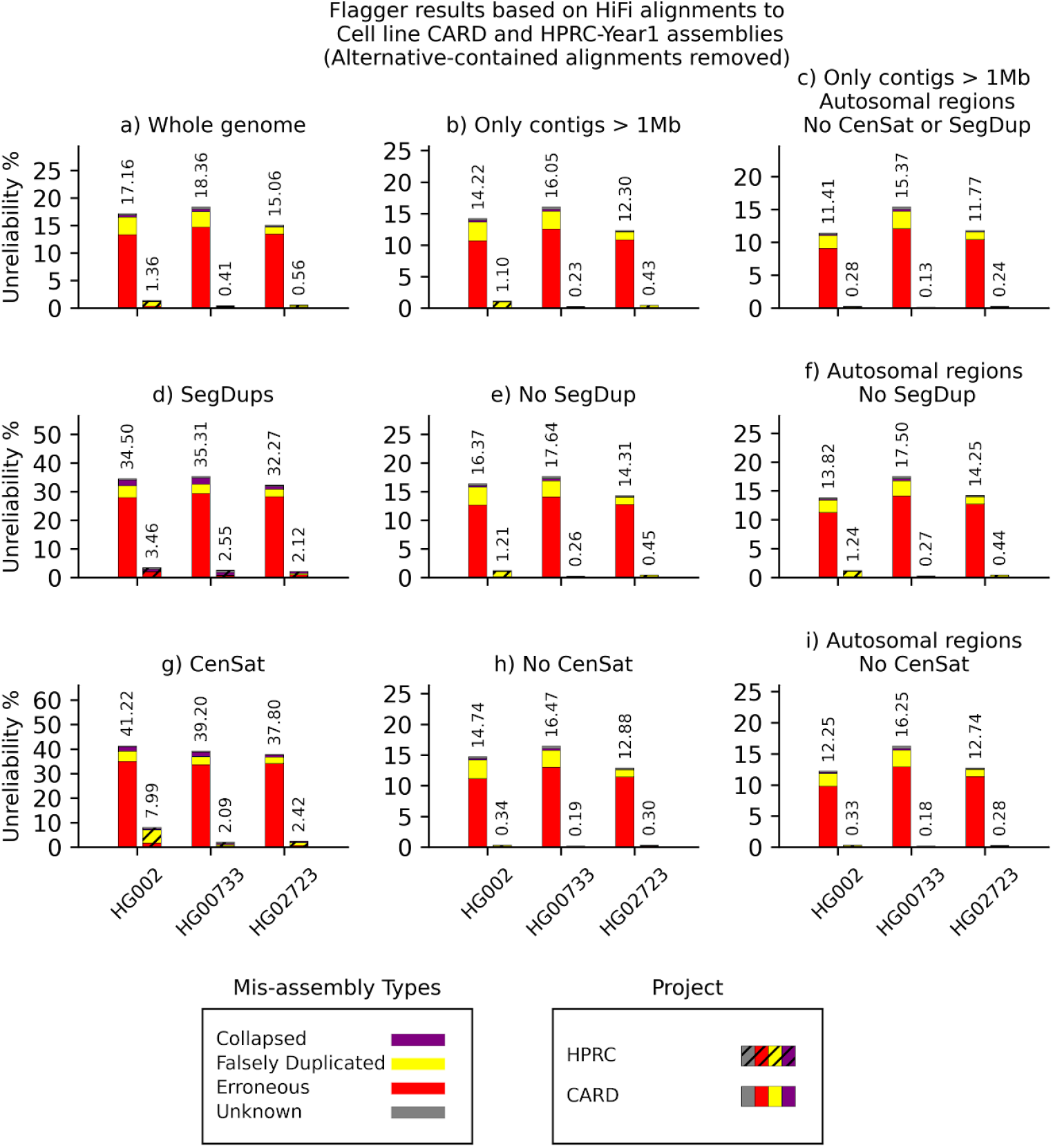
Flagger results based on ONT alignments to cell line CARD and HPRC-Y1 assemblies after removing alternative-contained alignments. The y axis of each panel indicates the unreliability percentages which are the total number of bases flagged as misassembly divided by the total assembly length and multiplied by one hundred

## References

100,000 Genomes Project Pilot Investigators, Smedley, D., Smith, K. R., Martin, A., Thomas, E. A., McDonagh, E. M., Cipriani, V., Ellingford, J. M., Arno, G., Tucci, A., Vandrovcova, J., Chan, G., Williams, H. J., Ratnaike, T., Wei, W., Stirrups, K., Ibanez, K., Moutsianas, L., Wielscher, M., … Caulfield, M. (2021). 100,000 Genomes Pilot on Rare-Disease Diagnosis in Health Care - Preliminary Report. The New England Journal of Medicine, 385(20), 1868–1880.

1000 Genomes Project Consortium, Abecasis, G. R., Auton, A., Brooks, L. D., DePristo, M. A., Durbin, R. M., Handsaker, R. E., Kang, H. M., Marth, G. T., & McVean, G. A. (2012). An integrated map of genetic variation from 1,092 human genomes. Nature, 491(7422), 56–65.

Cheng, H., Concepcion, G. T., Feng, X., Zhang, H., & Li, H. (2021). Haplotype-resolved de novo assembly using phased assembly graphs with hifiasm. Nature Methods, 18(2), 170–175.

Cheng, H., Jarvis, E. D., Fedrigo, O., Koepfli, K.-P., Urban, L., Gemmell, N. J., & Li, H. (2022). Haplotype-resolved assembly of diploid genomes without parental data. Nature Biotechnology, 40(9), 1332–1335.

Chen, X., Schulz-Trieglaff, O., Shaw, R., Barnes, B., Schlesinger, F., Källberg, M., Cox, A. J., Kruglyak, S., & Saunders, C. T. (2016). Manta: rapid detection of structural variants and indels for germline and cancer sequencing applications. Bioinformatics, 32(8), 1220–1222.

Chowdhury, M., Pedersen, B. S., Sedlazeck, F. J., Quinlan, A. R., & Layer, R. M. (2022). Searching thousands of genomes to classify somatic and novel structural variants using STIX. Nature Methods, 19(4), 445–448.

DePristo, M. A., Banks, E., Poplin, R., Garimella, K. V., Maguire, J. R., Hartl, C., Philippakis, A. A., del Angel, G., Rivas, M. A., Hanna, M., McKenna, A., Fennell, T. J., Kernytsky, A. M., Sivachenko, A. Y., Cibulskis, K., Gabriel, S. B., Altshuler, D., & Daly, M. J. (2011). A framework for variation discovery and genotyping using next-generation DNA sequencing data. Nature Genetics, 43(5), 491–498.

Ebler, J., Ebert, P., Clarke, W. E., Rausch, T., Audano, P. A., Houwaart, T., Mao, Y., Korbel, J. O., Eichler, E. E., Zody, M. C., Dilthey, A. T., & Marschall, T. (2022). Pangenome-based genome inference allows efficient and accurate genotyping across a wide spectrum of variant classes. Nature Genetics, 54(4), 518–525.

English, A. C., Menon, V. K., Gibbs, R. A., Metcalf, G. A., & Sedlazeck, F. J. (2022). Truvari: refined structural variant comparison preserves allelic diversity. Genome Biology, 23(1), 271.

Gibbs, J. R., van der Brug, M. P., Hernandez, D. G., Traynor, B. J., Nalls, M. A., Lai, S.-L., Arepalli, S., Dillman, A., Rafferty, I. P., Troncoso, J., Johnson, R., Zielke, H. R., Ferrucci, L., Longo, D. L., Cookson, M. R., & Singleton, A. B. (2010). Abundant quantitative trait loci exist for DNA methylation and gene expression in human brain. PLoS Genetics, 6(5), e1000952.

Heller, D., & Vingron, M. (2020). SVIM-asm: Structural variant detection from haploid and diploid genome assemblies. Bioinformatics, 36(22-23), 5519–5521.

Huang, K.-L., Mashl, R. J., Wu, Y., Ritter, D. I., Wang, J., Oh, C., Paczkowska, M., Reynolds, S., Wyczalkowski, M. A., Oak, N., Scott, A. D., Krassowski, M., Cherniack, A. D., Houlahan, K. E., Jayasinghe, R., Wang, L.-B., Zhou, D. C., Liu, D., Cao, S., … Ding, L. (2018). Pathogenic Germline Variants in 10,389 Adult Cancers. Cell, 173(2), 355–370.e14.

ICGC/TCGA Pan-Cancer Analysis of Whole Genomes Consortium. (2020). Pan-cancer analysis of whole genomes. Nature, 578(7793), 82–93.

Jain, M., Koren, S., Miga, K. H., Quick, J., Rand, A. C., Sasani, T. A., Tyson, J. R., Beggs, A. D., Dilthey, A. T., Fiddes, I. T., Malla, S., Marriott, H., Nieto, T., O’Grady, J., Olsen, H. E., Pedersen, B. S., Rhie, A., Richardson, H., Quinlan, A. R., … Loose, M. (2018). Nanopore sequencing and assembly of a human genome with ultra-long reads. Nature Biotechnology, 36(4), 338–345.

Jarvis, E. D., Formenti, G., Rhie, A., Guarracino, A., Yang, C., Wood, J., Tracey, A., Thibaud-Nissen, F., Vollger, M. R., Porubsky, D., Cheng, H., Asri, M., Logsdon, G. A., Carnevali, P., Chaisson, M. J. P., Chin, C.-S., Cody, S., Collins, J., Ebert, P., … Human Pangenome Reference Consortium. (2022). Automated assembly of high-quality diploid human reference genomes. In bioRxiv (p. 2022.03.06.483034). https://doi.org/10.1101/2022.03.06.483034

J Billingsley, K. (2022). Processing frozen human blood samples for population-scale Oxford Nanopore long-read DNA sequencing SOP v1. https://doi.org/10.17504/protocols.io.ewov1n93ygr2/v1

J Billingsley, K., Dewan, R., Malik, L., Alvarez Jerez, P., Kiley, S., Blauwendraat, C., & on behalf of the CARD Long-read Team. (2022). Processing human frontal cortex brain tissue for population-scale Oxford Nanopore long-read DNA sequencing SOP v2. https://doi.org/10.17504/protocols.io.kxygxzmmov8j/v2

Jiang, T., Liu, Y., Jiang, Y., Li, J., Gao, Y., Cui, Z., Liu, Y., Liu, B., & Wang, Y. (2020). Long-read-based human genomic structural variation detection with cuteSV. Genome Biology, 21(1), 189.

Karczewski, K. J., Francioli, L. C., Tiao, G., Cummings, B. B., Alföldi, J., Wang, Q., Collins, R. L., Laricchia, K. M., Ganna, A., Birnbaum, D. P., Gauthier, L. D., Brand, H., Solomonson, M., Watts, N. A., Rhodes, D., Singer-Berk, M., England, E. M., Seaby, E. G., Kosmicki, J. A., … MacArthur, D. G. (2020). The mutational constraint spectrum quantified from variation in 141,456 humans. Nature, 581(7809), 434–443.

Kirsche, M., Prabhu, G., Sherman, R., Ni, B., Aganezov, S., & Schatz, M. C. (2021). Jasmine: Population-scale structural variant comparison and analysis. In bioRxiv (p. 2021.05.27.445886). https://doi.org/10.1101/2021.05.27.445886

Kolmogorov, M., Yuan, J., Lin, Y., & Pevzner, P. A. (2019). Assembly of long, error-prone reads using repeat graphs. Nature Biotechnology, 37(5), 540–546.

Koren, S., Walenz, B. P., Berlin, K., Miller, J. R., Bergman, N. H., & Phillippy, A. M. (2017). Canu: scalable and accurate long-read assembly via adaptive -mer weighting and repeat separation. Genome Research, 27(5), 722–736.

Lee, H., & Schatz, M. C. (2012). Genomic dark matter: the reliability of short read mapping illustrated by the genome mappability score. Bioinformatics, 28(16), 2097–2105.

Liao, W.-W., Asri, M., Ebler, J., Doerr, D., Haukness, M., Hickey, G., Lu, S., Lucas, J. K., Monlong, J., Abel, H. J., Buonaiuto, S., Chang, X. H., Cheng, H., Chu, J., Colonna, V., Eizenga, J. M., Feng, X., Fischer, C., Fulton, R. S., … Paten, B. (2022). A Draft Human Pangenome Reference. In bioRxiv (p. 2022.07.09.499321). https://doi.org/10.1101/2022.07.09.499321

Li, H. (2018). Minimap2: pairwise alignment for nucleotide sequences. Bioinformatics, 34(18), 3094–3100.

Lin, J.-H., Chen, L.-C., Yu, S.-C., & Huang, Y.-T. (2022). LongPhase: an ultra-fast chromosome-scale phasing algorithm for small and large variants. Bioinformatics . https://doi.org/10.1093/bioinformatics/btac058

Lin, Y., Yuan, J., Kolmogorov, M., Shen, M. W., Chaisson, M., & Pevzner, P. A. (2016). Assembly of long error-prone reads using de Bruijn graphs. Proceedings of the National Academy of Sciences of the United States of America, 113(52), E8396–E8405.

Logsdon, G. A., Vollger, M. R., & Eichler, E. E. (2020). Long-read human genome sequencing and its applications. Nature Reviews. Genetics, 21(10), 597–614.

Loh, P.-R., Danecek, P., Palamara, P. F., Fuchsberger, C., A Reshef, Y., K Finucane, H., Schoenherr, S., Forer, L., McCarthy, S., Abecasis, G. R., Durbin, R., & L Price, A. (2016). Reference-based phasing using the Haplotype Reference Consortium panel. Nature Genetics, 48(11), 1443–1448.

Mahmoud, M., Doddapaneni, H., Timp, W., & Sedlazeck, F. J. (2021). PRINCESS: comprehensive detection of haplotype resolved SNVs, SVs, and methylation. Genome Biology, 22(1), 268.

Mahmoud, M., Gobet, N., Cruz-Dávalos, D. I., Mounier, N., Dessimoz, C., & Sedlazeck, F. J. (2019). Structural variant calling: the long and the short of it. Genome Biology, 20(1), 246.

Martin, M., Patterson, M., Garg, S., Fischer, S. O., Pisanti, N., Klau, G. W., Schöenhuth, A., & Marschall, T. (2016). WhatsHap: fast and accurate read-based phasing. In bioRxiv (p. 085050). https://doi.org/10.1101/085050

Mikheenko, A., Prjibelski, A., Saveliev, V., Antipov, D., & Gurevich, A. (2018). Versatile genome assembly evaluation with QUAST-LG. In Bioinformatics (Vol. 34, Issue 13, pp. i142–i150). https://doi.org/10.1093/bioinformatics/bty266

Nurk, S., Koren, S., Rhie, A., Rautiainen, M., Bzikadze, A. V., Mikheenko, A., Vollger, M. R., Altemose, N., Uralsky, L., Gershman, A., Aganezov, S., Hoyt, S. J., Diekhans, M., Logsdon, G. A., Alonge, M., Antonarakis, S. E., Borchers, M., Bouffard, G. G., Brooks, S. Y., … Phillippy, A. M. (2022). The complete sequence of a human genome. Science, 376(6588), 44–53.

Olson, N. D., Wagner, J., McDaniel, J., Stephens, S. H., Westreich, S. T., Prasanna, A. G., Johanson, E., Boja, E., Maier, E. J., Serang, O., Jáspez, D., Lorenzo-Salazar, J. M., Muñoz-Barrera, A., Rubio-Rodríguez, L. A., Flores, C., Kyriakidis, K., Malousi, A., Shafin, K., Pesout, T., … Zook, J. M. (2022). PrecisionFDA Truth Challenge V2: Calling variants from short and long reads in difficult-to-map regions. Cell Genomics, 2(5). https://doi.org/10.1016/j.xgen.2022.100129

Rautiainen, M., Nurk, S., Walenz, B. P., Logsdon, G. A., Porubsky, D., Rhie, A., Eichler, E. E., Phillippy, A. M., & Koren, S. (2022). Verkko: telomere-to-telomere assembly of diploid chromosomes. In bioRxiv (p. 2022.06.24.497523). https://doi.org/10.1101/2022.06.24.497523

Razaghi, R., Hook, P. W., Ou, S., Schatz, M. C., Hansen, K. D., Jain, M., & Timp, W. (2022). Modbamtools: Analysis of single-molecule epigenetic data for long-range profiling, heterogeneity, and clustering. In bioRxiv (p. 2022.07.07.499188). https://doi.org/10.1101/2022.07.07.499188

Rhie, A., McCarthy, S. A., Fedrigo, O., Damas, J., Formenti, G., Koren, S., Uliano-Silva, M., Chow, W., Fungtammasan, A., Kim, J., Lee, C., Ko, B. J., Chaisson, M., Gedman, G. L., Cantin, L. J., Thibaud-Nissen, F., Haggerty, L., Bista, I., Smith, M., … Jarvis, E. D. (2021). Towards complete and error-free genome assemblies of all vertebrate species. Nature, 592(7856), 737–746.

Schatz, M. C., Philippakis, A. A., Afgan, E., Banks, E., Carey, V. J., Carroll, R. J., Culotti, A., Ellrott, K., Goecks, J., Grossman, R. L., Hall, I. M., Hansen, K. D., Lawson, J., Leek, J. T., Luria, A. O., Mosher, S., Morgan, M., Nekrutenko, A., O’Connor, B. D., … Wuichet, K. (2022). Inverting the model of genomics data sharing with the NHGRI Genomic Data Science Analysis, Visualization, and Informatics Lab-space. Cell Genomics, 2(1). https://doi.org/10.1016/j.xgen.2021.100085

Sedlazeck, F. J., Lee, H., Darby, C. A., & Schatz, M. C. (2018). Piercing the dark matter: bioinformatics of long-range sequencing and mapping. Nature Reviews. Genetics, 19(6), 329–346.

Sedlazeck, F. J., Rescheneder, P., Smolka, M., Fang, H., Nattestad, M., von Haeseler, A., & Schatz, M. C. (2018). Accurate detection of complex structural variations using single-molecule sequencing. Nature Methods, 15(6), 461–468.

Shafin, K., Pesout, T., Chang, P.-C., Nattestad, M., Kolesnikov, A., Goel, S., Baid, G., Kolmogorov, M., Eizenga, J. M., Miga, K. H., Carnevali, P., Jain, M., Carroll, A., & Paten, B. (2021). Haplotype-aware variant calling with PEPPER-Margin-DeepVariant enables high accuracy in nanopore long-reads. Nature Methods, 18(11), 1322–1332.

Shafin, K., Pesout, T., Lorig-Roach, R., Haukness, M., Olsen, H. E., Bosworth, C., Armstrong, J., Tigyi, K., Maurer, N., Koren, S., Sedlazeck, F. J., Marschall, T., Mayes, S., Costa, V., Zook, J. M., Liu, K. J., Kilburn, D., Sorensen, M., Munson, K. M., … Paten, B. (2020). Nanopore sequencing and the Shasta toolkit enable efficient de novo assembly of eleven human genomes. Nature Biotechnology, 38(9), 1044–1053.

Sirén, J., Monlong, J., Chang, X., Novak, A. M., Eizenga, J. M., Markello, C., Sibbesen, J. A., Hickey, G., Chang, P.-C., Carroll, A., Gupta, N., Gabriel, S., Blackwell, T. W., Ratan, A., Taylor, K. D., Rich, S. S., Rotter, J. I., Haussler, D., Garrison, E., & Paten, B. (2021). Pangenomics enables genotyping of known structural variants in 5202 diverse genomes. Science, 374(6574), abg8871.

Smolka, M., Paulin, L. F., Grochowski, C. M., Mahmoud, M., Behera, S., Gandhi, M., Hong, K., Pehlivan, D., Scholz, S. W., Carvalho, C. M. B., Proukakis, C., & Sedlazeck, F. J. (2022). Comprehensive Structural Variant Detection: From Mosaic to Population-Level. In bioRxiv (p. 2022.04.04.487055). https://doi.org/10.1101/2022.04.04.487055

Vollger, M. R., Dishuck, P. C., Sorensen, M., Welch, A. E., Dang, V., Dougherty, M. L., Graves-Lindsay, T. A., Wilson, R. K., Chaisson, M. J. P., & Eichler, E. E. (2019). Long-read sequence and assembly of segmental duplications. Nature Methods, 16(1), 88–94.

Wagner, J., Olson, N. D., Harris, L., Khan, Z., Farek, J., Mahmoud, M., Stankovic, A., Kovacevic, V., Yoo, B., Miller, N., Rosenfeld, J. A., Ni, B., Zarate, S., Kirsche, M., Aganezov, S., Schatz, M. C., Narzisi, G., Byrska-Bishop, M., Clarke, W., … Zook, J. M. (2022). Benchmarking challenging small variants with linked and long reads. Cell Genomics, 2(5). https://doi.org/10.1016/j.xgen.2022.100128

Wagner, J., Olson, N. D., Harris, L., McDaniel, J., Cheng, H., Fungtammasan, A., Hwang, Y.-C., Gupta, R., Wenger, A. M., Rowell, W. J., Khan, Z. M., Farek, J., Zhu, Y., Pisupati, A., Mahmoud, M., Xiao, C., Yoo, B., Sahraeian, S. M. E., Miller, D. E., … Sedlazeck, F. J. (2022). Curated variation benchmarks for challenging medically relevant autosomal genes. Nature Biotechnology, 40(5), 672–680.

Zarate, S., Carroll, A., Mahmoud, M., Krasheninina, O., Jun, G., Salerno, W. J., Schatz, M. C., Boerwinkle, E., Gibbs, R. A., & Sedlazeck, F. J. (2020). Parliament2: Accurate structural variant calling at scale. GigaScience, 9(12). https://doi.org/10.1093/gigascience/giaa145

Zook, J. M., Hansen, N. F., Olson, N. D., Chapman, L., Mullikin, J. C., Xiao, C., Sherry, S., Koren, S., Phillippy, A. M., Boutros, P. C., Sahraeian, S. M. E., Huang, V., Rouette, A., Alexander, N., Mason, C. E., Hajirasouliha, I., Ricketts, C., Lee, J., Tearle, R., … Salit, M. (2020). A robust benchmark for detection of germline large deletions and insertions. Nature Biotechnology, 38(11), 1347–1355.

